# Mussels on the move: new records of the invasive non-native quagga mussel (*Dreissena rostriformis bugensis*) in Great Britain using eDNA and a new probe-based qPCR assay

**DOI:** 10.1101/2023.12.18.572119

**Authors:** Sara Peixoto, Rosetta C. Blackman, Jonathan Porter, Alan Wan, Chris Gerrard, Ben Aston, Lori Lawson Handley

## Abstract

Invasive non-native species (INNS) pose a worldwide environmental threat, negatively impacting invaded ecosystems on an ecological and economical scale. In recent decades, quagga mussels (*Dreissena rostriformis bugensis*) have successfully invaded several countries in Western Europe from the Ponto-Caspian region, being recorded for the first time in Great Britain (GB) in 2014, in Wraysbury, near London. In recent years, environmental DNA (eDNA) analysis has proven to be a sensitive and effective method for early detection and monitoring of a number of INNS. Previously, a dye-based quantitative PCR (qPCR) assay was developed for the detection of quagga mussels from eDNA samples. Here, a target-specific probe was designed to further increase the specificity of this assay and used to obtain an updated distribution of this species in GB. Twenty-four sites were sampled, including sites with established populations near London and sites spread across the East Midlands and East Anglia regions. Positive detections were obtained for 11 of the 24 sites, and these were widely spread, as far as Nottingham (East Midlands) and Norfolk (East Anglia). Detection rates were 100% at the three sites with known established populations, while rates were lower (3-50% of positive replicates) in the eight newly-identified sites, consistent with an early stage of invasion. Of particular concern was the detection of quagga mussels in major waterways and in popular recreational sites, highlighting urgent measures are needed to control pathways and spread. Our study demonstrates that quagga mussels are considerably more widespread in GB than previously thought and provides a much-needed step towards operational use of eDNA for monitoring quagga mussels.

## Introduction

Invasive non-native species (INNS) pose a worldwide environmental threat, causing ecological and economic impacts on invaded ecosystems. Ponto-Caspian invaders such as the quagga mussel (*Dreissena rostriformis bugensis*, Andrusov, 1897) are of special concern due their successful large-scale invasion into Western Europe in recent decades. The first observation of quagga mussels in Western Europe dates back to 2006 in The Netherlands (Molloy et al., 2007). This introduction was suggested to be either via the Main-Danube Canal (Molloy et al., 2007) or due to the discharge of contaminated ballast water in the port of Rotterdam (Velde & Platvoet, 2007). River connectivity within Europe allowed for further spread, and in the following years more records of quagga mussels were documented, with first detections in Germany in 2007 (Velde & Platvoet, 2007), Belgium in 2010 (Marescaux & Van Doninck, 2012), France in 2011 (Bij de Vaate & Beisel, 2011), Switzerland in 2015 (Haltiner et al., 2022), and Italy in 2022 (Salmaso et al., 2022).

In England, the first quagga mussel record was documented during routine monitoring in 2014 in Wraysbury River, a tributary of the River Thames, near London (Aldridge, Ho & Froufe, 2014). This observation occurred shortly after a horizon scanning study identified quagga mussels as the non-native species with the highest risk of invasion, establishment, and impacts in Great Britain (GB), posing major threats to Britain’s freshwater biodiversity (Roy et al., 2014). After almost a decade, quagga mussel populations are now well established in the Thames and its tributaries. With the River Thames serving as a major corridor to the wider canal and river network (Aldridge et al., 2014), populations of quagga mussels have since been discovered in reservoirs and other waterways north of London (National Biodiversity Network Trust, 2023; Fig. 1). More recently, they have been found further north in Rutland Water reservoir (Environment Agency, 2020) and in a water treatment facility in Lincoln (Aldridge, 2023; Fig. 1), making the latter the northernmost point they have been recorded so far.

**Fig. 1.**
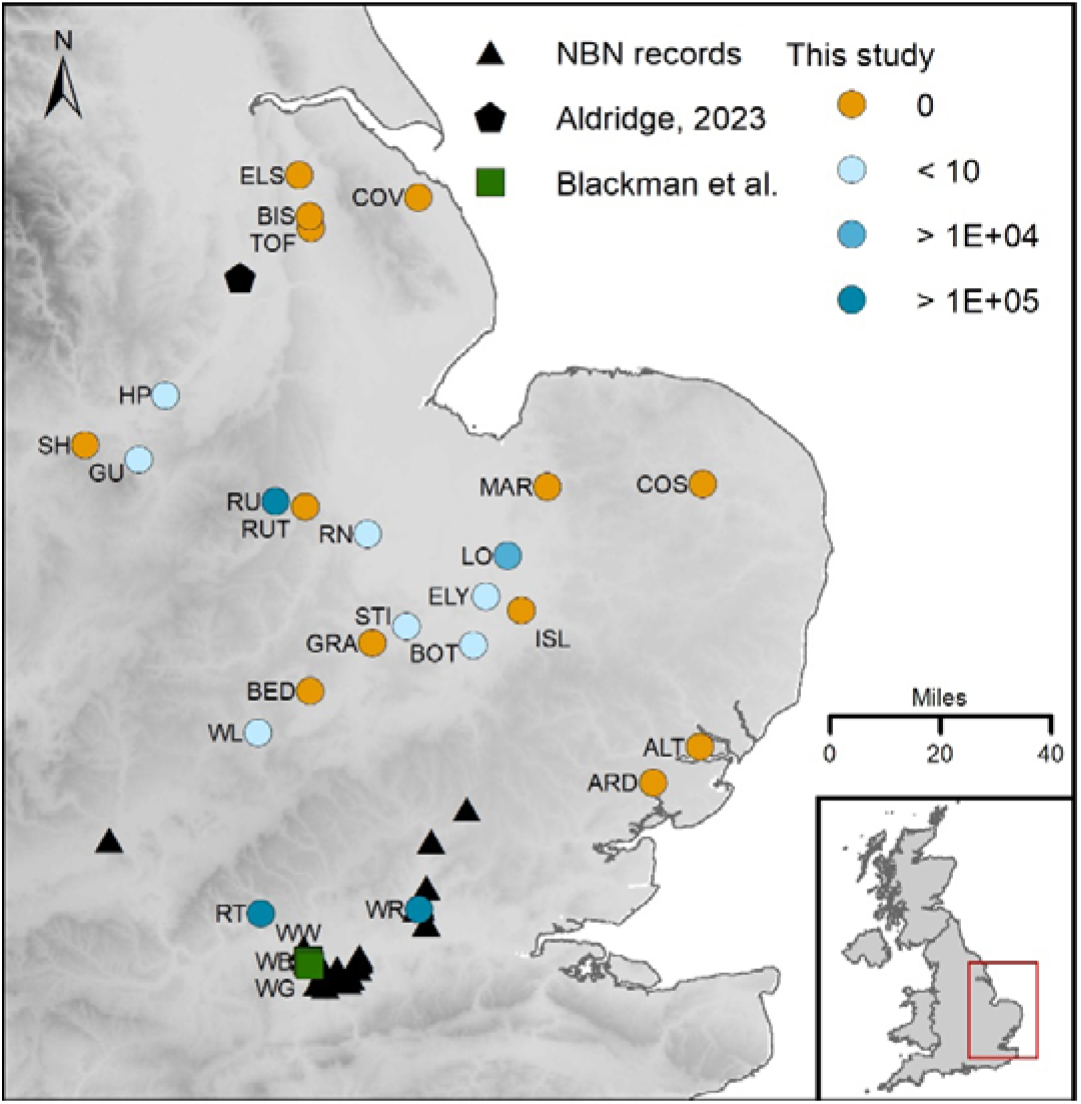
Location of the 24 sites sampled in England for this study (circles) and respective average DNA copies/μL recovered. Green squares represent the locations of the eDNA samples collected by Blackman et al. (2020b) and used here for assay optimization. Site codes are the same as in Table 2. Black data points correspond to the current known distribution of quagga mussels, obtained from the National Biodiversity Network (NBN) Atlas, as of April 5^th^ 2023 (triangles) and from Aldridge (2023) (pentagon). They have also been recorded at Rutland Water (RU), however this is not depicted in the map due to overlap with data points from this study.

Species distribution models based on bioclimatic data predict that the distribution of quagga mussels could extend to much of England, central Scotland and southern Wales (Gallardo and Aldridge, 2013). It is often assumed that their distribution will be similar to that of zebra mussels, which have been established in GB for over 100 years and are widespread (Aldridge et al., 2014). Nevertheless, environmental models suggest that there are important differences in the two species’ niches, with quagga mussels preferring higher temperatures and lower precipitation (Quinn, Gallardo & Aldridge, 2014), and being more tolerant of low oxygen (Karatayev, Burlakova & Padilla, 1998). Moreover, while zebra mussels require hard substrates and are more abundant in the littoral zone, quagga mussels are also able to attach to and colonise soft substrates such as silty sediments, and favour the profundal zones of lakes (Karatayev, Burlakova & Padilla, 2015; Karatayev & Burlakova, 2022).

Quagga mussels are generally more competitive than zebra mussels, and in most sites where the two species co-occur they can quickly outcompete and displace zebra mussels (Haltiner et al., 2022; Karatayev et al., 2021; Strayer et al., 2019). A study in which larvae of both species were reared in controlled conditions showed that the planktonic stage of quagga mussels (veligers) took more time to settle (Wright et al., 1996), suggesting an extended presence in the water column which could allow them to disperse over longer distances. Moreover, quagga mussels are able to reproduce at lower temperatures than zebra mussels (Karatayev & Burlakova, 2022) and larvae are usually found year-round in invaded sites (e.g. Haltiner et al., 2022). Their ability to survive and grow at lower temperatures and with less food is reflected in greater ecological impacts on invaded ecosystems and for longer periods of time (Karatayev & Burlakova, 2022).

Both dreissenid species are ecosystem engineers due to their high water filtering capacity, which has important direct and indirect effects on invaded systems (MacIsaac, 1996; Roy et al., 2014). Quagga mussels quickly proliferate to become dominant, causing ecological impacts that range from changes to the density and richness of macroinvertebrate communities (Mills, Chadwick & Francis, 2019; Ward and Ricciardi, 2007), fouling and suffocation of unionid mussels (Larson, Bailey & Evans, 2022; Schloesser et al., 2006), and modifications of the river’s geomorphic processes such as sediment mobility (Sanders et al., 2022). In addition to ecological impacts, the biofouling of infrastructures on invaded sites poses a problem, particularly for water companies (Chakraborti et al., 2013). The presence of quagga mussels on infested water treatment plants requires increased maintenance to keep the components (e.g., pipes, tanks, intake structure) clean and this is associated with elevated costs (Chakraborti, Madon & Kaur, 2016; Connelly et al., 2007). The potential for spread and high impact, both ecologically and economically, means that the quagga mussel is recognised as a top priority species for monitoring and mitigation (Roy et al., 2014).

Sensitive tools that allow early detection and rapid response are crucial for monitoring invasive species such as quagga mussels. The analysis of environmental DNA (eDNA) samples (e.g. soil, water, air) has proven to be an efficient method for the detection of INNS from different taxonomic groups and different environments, often outperforming traditional methods (Blackman, Hänfling & Lawson-Handley, 2018; Fonseca et al., 2023; Lawson Handley, 2015). Likewise, quantitative PCR (qPCR) has proven to be an efficient and sensitive method and has been the technique of choice over the years for the detection of several invasive species, across different taxonomic groups (e.g. Gingera et al., 2017; Prabhakaran et al., 2023; Roux et al., 2020).

Several targeted eDNA assays already exist for the detection of dreissenid mussels from eDNA samples (reviewed in Feist & Lance, 2021), however some of these co-amplify both quagga and zebra mussels (e.g. Gingera et al., 2017; Peñarrubia et al., 2016). Recently, sensitive and species-specific conventional PCR and dye-based qPCR assays (“DRB1”) were developed for quagga mussels and tested *in silico, in vitro*, in mesocosm experiments and field trials (Blackman et al., 2020a, 2020b). Both the conventional and dye-based qPCR assays, from here referred to as cDRB1 and dDRB1 respectively, outperformed kick-sampling and eDNA metabarcoding for the detection of quagga mussels in field trials conducted in the Wraysbury River, and the qPCR assay had the advantage of providing information on the decreasing signal of DNA concentration with increasing distance from the main source population (Blackman et al., 2020b).

Understanding the uncertainties and limitations associated with targeted eDNA assays, such as the rate of false positives and negatives, helps policymakers and end-users to choose the best assay for routine monitoring. With this in mind, a 5-level validation scale for targeted eDNA assays was developed by Thalinger et al. (2021), with minimum criteria defined for each level, ranging from level 1 (“incomplete”) to level 5 (“operational”). The dDRB1 assay currently meets the minimum requirements of levels 1-3, i.e., *in silico* analysis, *in vitro* testing on target tissue and closely related species, and detection from environmental samples (Blackman et al., 2020a, 2020b). However, it is well recognised that inclusion of a target-specific probe during qPCR increases assay specificity and reduces the chance of false-positives, providing more confidence in the results (Thalinger et al., 2021). Further development and testing of the DRB1 assay is therefore required to improve the readiness of the assay for routine monitoring. The goals of this study were thus to 1) further improve the dDRB1 assay, by designing a probe and estimating limits of detection (LOD), and 2) use the probe-based qPCR assay (from here referred to as pDRB1) to screen for quagga mussels in several locations in GB, in order to update their current distribution.

## Materials and methods

### Development of pDRB1 assay

The quagga mussel-specific DRB1 assay (Blackman et al., 2020a, 2020b), targeting a 188 bp fragment of the cytochrome oxidase I (COI) gene, was further developed by designing a target-specific probe (Table 1). A TaqMan probe was designed using the PrimerQuest tool (IDT, www.idtdna.com) in conjunction with alignments (Clustal Omega) of quagga and related mussel COI sequences from the EBI database. Candidate probe sequences were confirmed *in silico* using the EBI database with the consensus target sequences used for eventual quantification standards (Table 1).

**Table 1.**
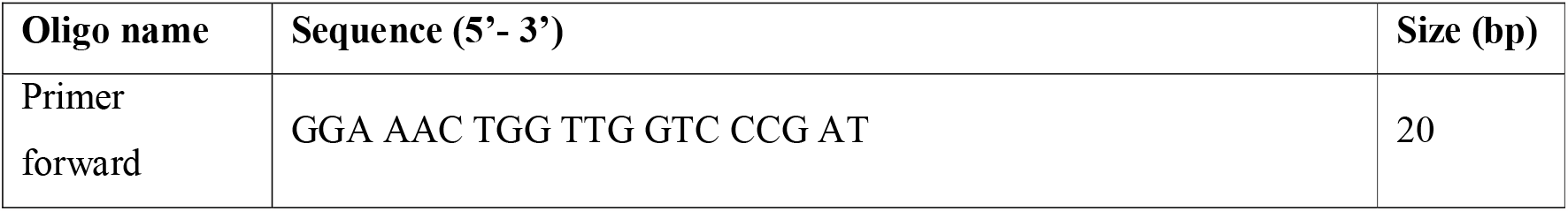
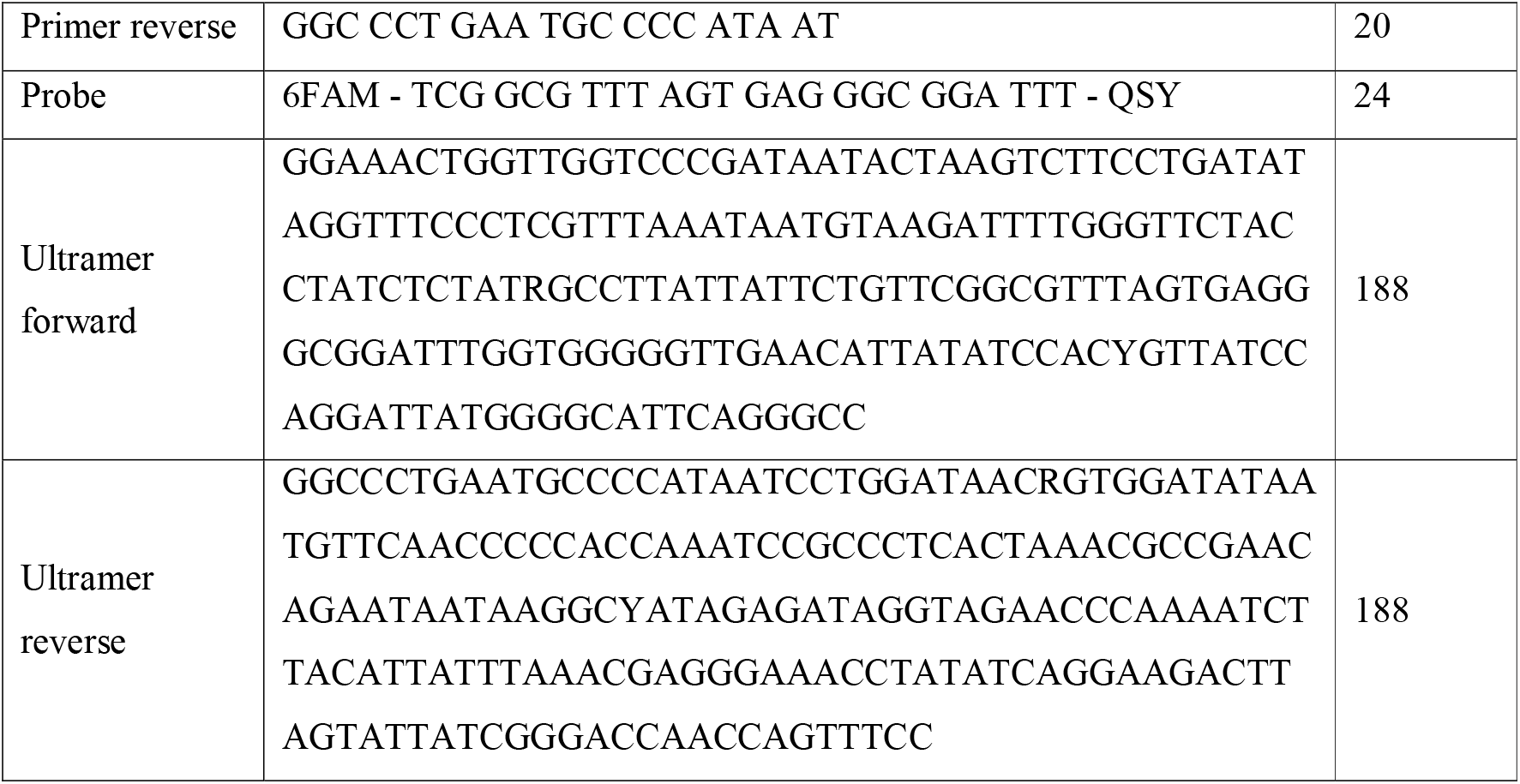
Details of the species-specific pDRB1 assay, including primers, probe, and ultramers (used for the standard curve).

The new primer and probe combination was then optimised *in vitro* by varying annealing temperatures against tissue of quagga mussels, closely related taxa - zebra mussel (*Dreissena polymorpha*), killer shrimp (*Dikerogammarus villosus*) and demon shrimp (*Dikerogammarus haemobaphes*) - and common species found in GB - pea mussel (*Sphaerium corneum*), blue mussel (*Mytilus edulis*), European oyster (*Ostrea edulis*), common periwinkle (*Littorina littorea*) and the common limpet (*Patella vulgata*). Standards made of the target amplicon, which included the primer and probe binding sites (ultramers; Table 1), and with known copy numbers were used to assess the assay’s efficiency. For this, four 10-fold dilutions ranging from 10^1^ to 10^4^ copies/μL numbers were run together with the tissue samples. Each standard was run twice and tissue samples four times. qPCR reactions were performed with 12.5 μL of TaqMan Universal PCR Master Mix (Fisher Scientific, UK), 1.6 μM of primers forward and reverse combined, 0.05 μM of probe, 0.64 mg/ml of BSA and 2 μL of sample. The qPCR thermal profile consisted of an initial step at 50°C for 10 min and 95°C for 10 min, followed by 40 cycles of 95°C for 15 seconds and annealing for 1 min at 60, 62 and 63°C.

eDNA water samples from Wraysbury River, previously collected and tested with the dDRB1 assay by Blackman et al. (2020b), were tested with the new probe-based assay to compare the performance of both assays. Three samples were selected from three different locations - Wraysbury Bridge, Wraysbury Gardens, Wraysbury Weir. A total of nine samples were chosen based on DNA copy numbers, in order to include a range of low, medium and high DNA concentrations. Final qPCR conditions used for eDNA samples were as described above but with 45 cycles and 62°C as the annealing temperature. Six replicates were performed for each sample.

### eDNA sample collection

Sample collection was conducted in England, with a focus on the area between the River Thames (at Bourne End, Buckinghamshire) in the south, to the River Ancholme (Lincolnshire) in the north, and from Staunton Harold Reservoir (Derbyshire) in the west to Norfolk and Suffolk in the east (Fig. 1). Sampling locations were chosen to include known established populations of quagga mussels (i.e. Rutland Water, Walthamstow Reservoirs, River Thames), priority sites with suspected but unconfirmed reports (i.e. Holme Pierrepont water park and Grand Union canal), sites upstream of water company intakes and sites adjacent to already invaded water bodies, while providing a broad coverage across the East Midlands and East Anglian regions. Water samples were collected from a total of 24 sites including 1 canal, 2 lakes, 5 reservoirs and 16 rivers (Fig. 1; Table 2). Five water samples were collected at each site, with the exception of Ardleigh (ARD) where only three samples were collected due to lack of access. Samples were collected from five locations spaced around 100 m, although this was not always possible due to safety reasons, and in such cases samples were collected within smaller distances or in the same location (see Table S1 for coordinates for each individual sample). Water was collected using a sterile whirl pak bag, without disturbing the sediment. Gloves were worn at all times during handling of samples to minimise contamination and new sterile equipment was used for each sample. A field negative control of purified water brought from the lab or shop-bought water was filtered at each site.

**Table 2.**
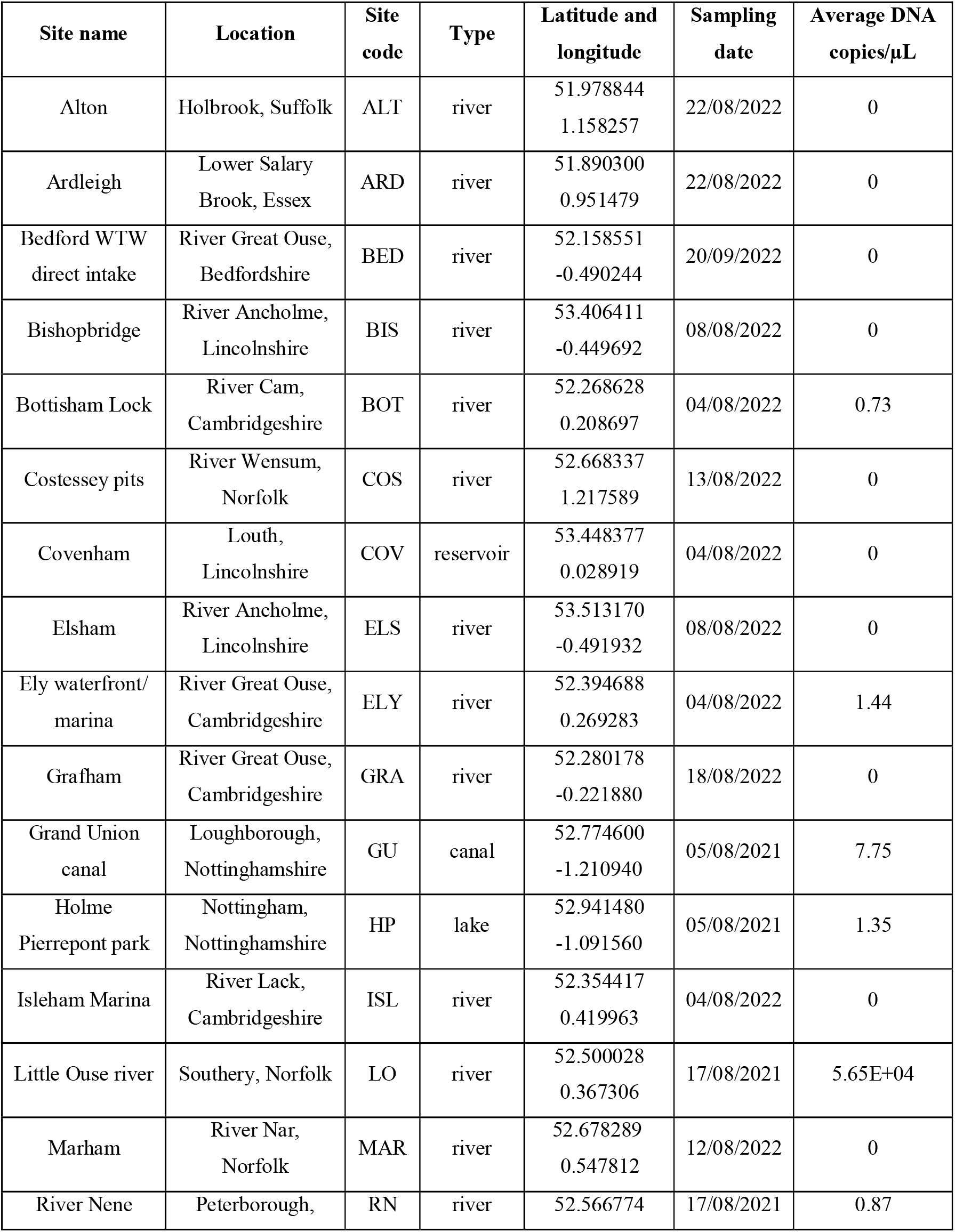
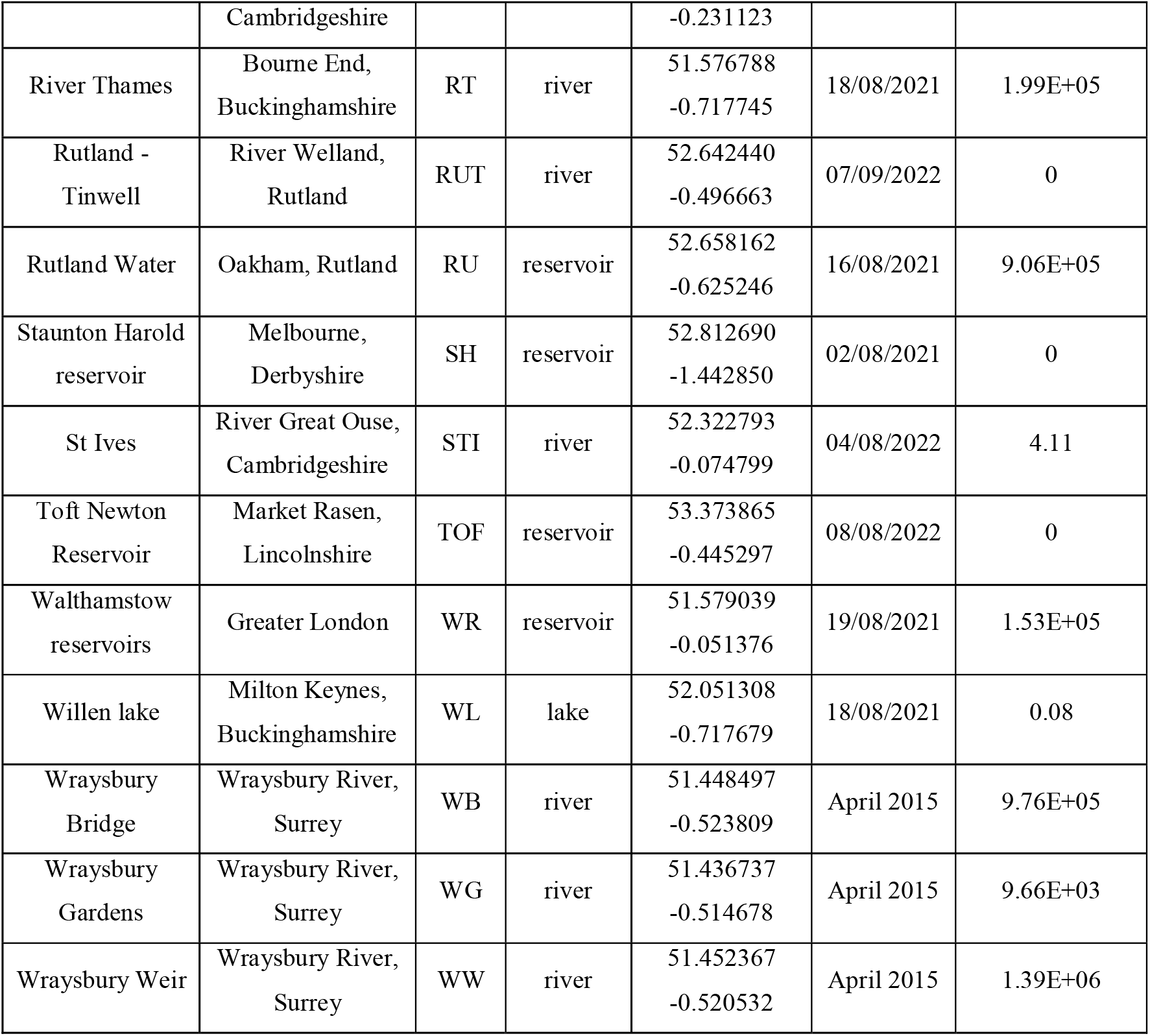
Details of the 24 locations sampled in this study as well as the three sites at Wraysbury River (sampled by Blackman et al., 2020b) that were used for testing the new pDRB1 assay. Sites are ordered alphabetically.

For nine of the 24 sites, sample collection was conducted in August 2021 (Table 2). Water samples were filtered using a 100 ml luer lock syringe (Nature Metrics, UK) and an enclosed filter with a polyethersulfone membrane and 0.8 μm pore size (Nature Metrics, UK). Water was pushed through the filter as many times as possible until the filter clogged, and the volume filtered was recorded (Table S1). Air was passed through the filter to dry it and 1 ml of Longmire’s buffer was added to each filter to preserve the sample. Following sample collection, filters were stored at room temperature until return to the laboratory, where they were stored at -20°C until DNA extraction. For the remaining 15 sites, sample collection was conducted during August and September 2022 (Table 2). Water was collected and preserved as previously described, but in this case, due to unavailability of the filters formerly used, filtration was conducted using Sterivex filters with 0.45 μm pore size and a PVDF (polyvinylidene difluoride) membrane (Merck Millipore, UK) and a 60 ml luer lock syringe (Fisher Scientific, UK).

### DNA extraction

Samples collected with Nature Metrics filters were extracted using a modified version of the DNeasy Blood & Tissue Kit (Qiagen, UK), whereas samples obtained with Sterivex filters were extracted using a modified version of the Mu-DNA protocol (Di Muri et al., 2020; Sellers et al., 2018). Both protocols are available in the supplementary material. All DNA extractions were conducted in a dedicated laboratory for processing eDNA samples and a negative control was included in each extraction batch to monitor for contamination. Following extractions, the purity and concentration of all samples were assessed using a Nanodrop 1000 spectrophotometer (Thermo Fisher Scientific, UK).

### Inhibition tests

All samples used in this study (i.e. samples collected in the present study and samples from Blackman et al., 2020b) were tested for inhibition prior to qPCR with the pDRB1 assay, using the Applied Biosystems TaqMan Exogenous Internal Positive Control Reagents (Fisher Scientific, UK). A similar protocol was applied for all samples with the only difference being the type of master mix used (Table S1). For samples collected with Nature Metrics filters and extracted with the Qiagen kit, the TaqMan Environmental Master Mix 2.0 (Fisher Scientific, UK) was used, while for the remaining samples the TaqMan Universal Master Mix 2.0 (Fisher Scientific, UK) was used instead. This was due to the inefficiency of dilution in overcoming inhibition for the first set of samples when using the TaqMan Universal Master Mix 2.0, which was only overcome when using the TaqMan Environmental Master Mix 2.0.

Reaction volumes consisted of 7.5 μL of TaqMan Environmental or Universal Master Mix 2.0 (Fisher Scientific, UK), 1.5 μL of 10X Exo IPC Mix, 0.3 μL of 50X Exo IPC DNA, 3.7 μL of molecular grade water and 2 μL of sample. Reactions were run on a StepOnePlus Real-Time PCR machine with the following thermal cycler conditions: 2 min at 50°C and 10 min at 95°C followed by 40 cycles of 15 s at 95°C and 1 min at 60°C. All eDNA samples were tested in duplicate and samples were considered to be inhibited if the average cycle threshold (Cq) of a sample was higher than the no template reaction by 2 or more cycles (e.g. Tillotson et al., 2018). All samples that showed inhibition were diluted 10x and re-run with the respective master mix to confirm whether inhibition was overcome (Table S1). If amplification with the species-specific qPCR assay was detected for diluted samples, a 10x correction factor was applied to their final DNA concentration.

### eDNA sample screening

Following inhibition tests, eDNA samples were analysed with the newly developed pDRB1 assay. Final qPCR conditions for eDNA samples consisted of 12.5 μL of TaqMan Environmental or Universal Master Mix 2.0 (the same used for the inhibition step), 1.6 μM of primers (forward and reverse combined), 0.05 μM of probe (Table 1), 0.64 mg/ml of BSA, 5.45 μL of water and 2 μL of sample. The qPCR program consisted of an initial step at 50°C for 10 min and 95°C for 10 min, followed by 45 cycles of 95°C for 15 seconds and 62°C for 1 min on a StepOnePlus Real-Time PCR machine. Six replicates were performed for each eDNA sample and respective field and extraction negative control. Eight qPCR negative controls were included in each plate to monitor for contamination during PCR preparation.

Standards of known concentration were included in each plate, in triplicate, to accurately quantify the DNA concentration of eDNA samples. For this, two single stranded DNA sequences of 188 bp each (Table 1; Integrated DNA Technologies, Belgium) were combined and serially diluted to obtain four 10-fold dilutions ranging from 10^4^ to 10^1^ copies/μL.

### Data analyses

To compare the performance of the dDRB1 and pDRB1 assays, a non-parametric Wilcoxon signed-rank test was used. For this, the average DNA copies/μL for each of the nine samples collected by Blackman et al. (2020b) and each assay was calculated and tested with the “wilcox.test()” function from the stats package in R (R Core Team, 2022). The map was created with ArcMap 10.8.2 and the remaining figures were made using the ggplot2 R package (Wickham, 2016).

## Results

### Development of pDRB1 assay

All annealing temperatures tested showed species-specific amplification of quagga mussel with no cross-amplification of other species. qPCR amplification efficiency values were 102.9, 99.1 and 63.4% for annealing temperatures of 60, 62 and 63°C, respectively. At 63°C, the lowest standard (10^1^) did not show any amplification, which explains the lower efficiency value. Cq values for quagga mussel amplification between 60 and 62°C were very similar (21.65 and 21.61, respectively). All annealing temperatures showed little or lack of a plateau stage when run for 40 cycles only. The final retained conditions included 62°C as the annealing temperature and 45 cycles.

There was a statistically significant difference between the average DNA copies/μL obtained for dDRB1 (mean = 3.87+05, SD = 5.84+05) and pDRB1 (mean = 7.93+05, SD = 1.07+06) assays (Wilcoxon signed-rank test: V = 0, p = 0.004, n = 9), with the latter consistently yielding higher DNA copy numbers (Fig. 2). Details on Cq values and copies/μL for each sample and assay is provided in the supplementary material (Table S2).

**Fig. 2.**
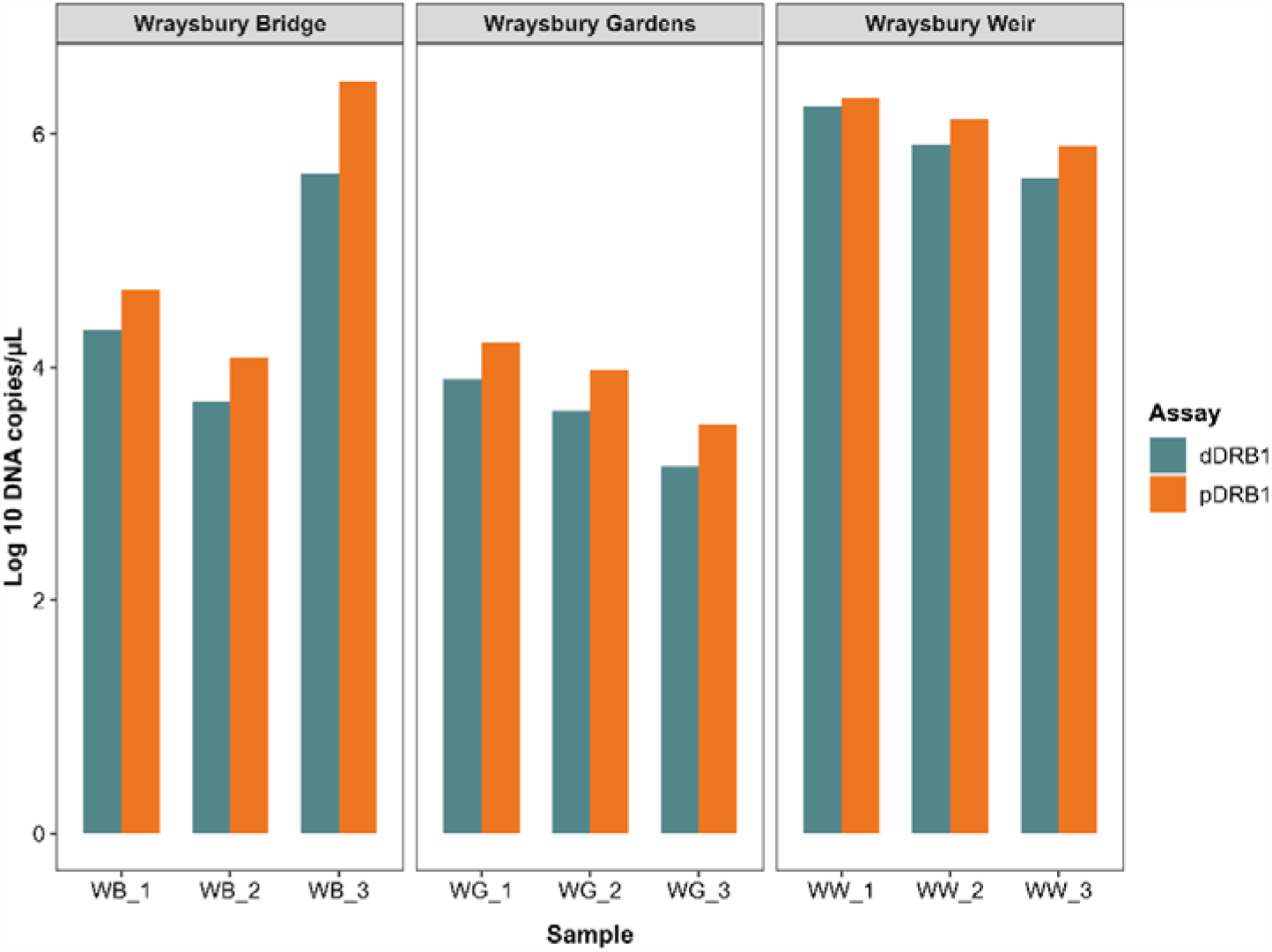
Number of log 10 DNA copies/μL obtained for both DRB1 assays for each of the 9 samples tested from Blackman et al. (2020b). For visual purposes log 10 DNA copies/μL is being used instead of non-transformed data.

Further information about the assay and MIQE checklist (Bustin et al., 2009) can be found in the supplementary material (Table S3).

### Inhibition tests

Out of the 118 samples from this study, inhibition was observed in 21 samples from nine sites (Table S1). In addition, one sample from the nine collected by Blackman et al. (2020b) showed inhibition. The 10x dilution was successful in overcoming inhibition for all the samples.

### eDNA sample screening

All standards amplified in 100% of the replicates with the exception of the lowest one (10 copies/μL) which amplified in 93% of the replicates. The LOD, i.e. the lowest standard with at least 95% amplification, was therefore 100 copies/μL. qPCR assays exhibited an average efficiency of 97.2% (91.8 – 104.8) and R^2^ of 0.993 (0.984-0.998).

From the 24 sites sampled in our study, 11 sites had detectable quagga mussel DNA (Fig. 1 and 3). All samples and replicates amplified for three sites - Rutland Water, Walthamstow Reservoirs and River Thames - with average DNA copy numbers ranging from 1.5E+05 to 9.1E+05 copies/μL (Fig. 1; Table 2). Two further sites were positive for all 5 samples - Grand Union canal and Little Ouse river - although with only 15 and 14 positive replicates out of 30, respectively (Fig. 3). The additional six sites with detectable quagga mussel DNA exhibited very low detection levels, with a maximum of 3/5 (60%) samples and 7/30 (23%) of replicates amplifying (Fig. 3). No amplification was observed for any sample of the remaining 13 sites (Fig. 1). Information on Cq values and copies/μL for each sample and replicate is provided in the supplementary material (Table S4).

**Fig. 3.**
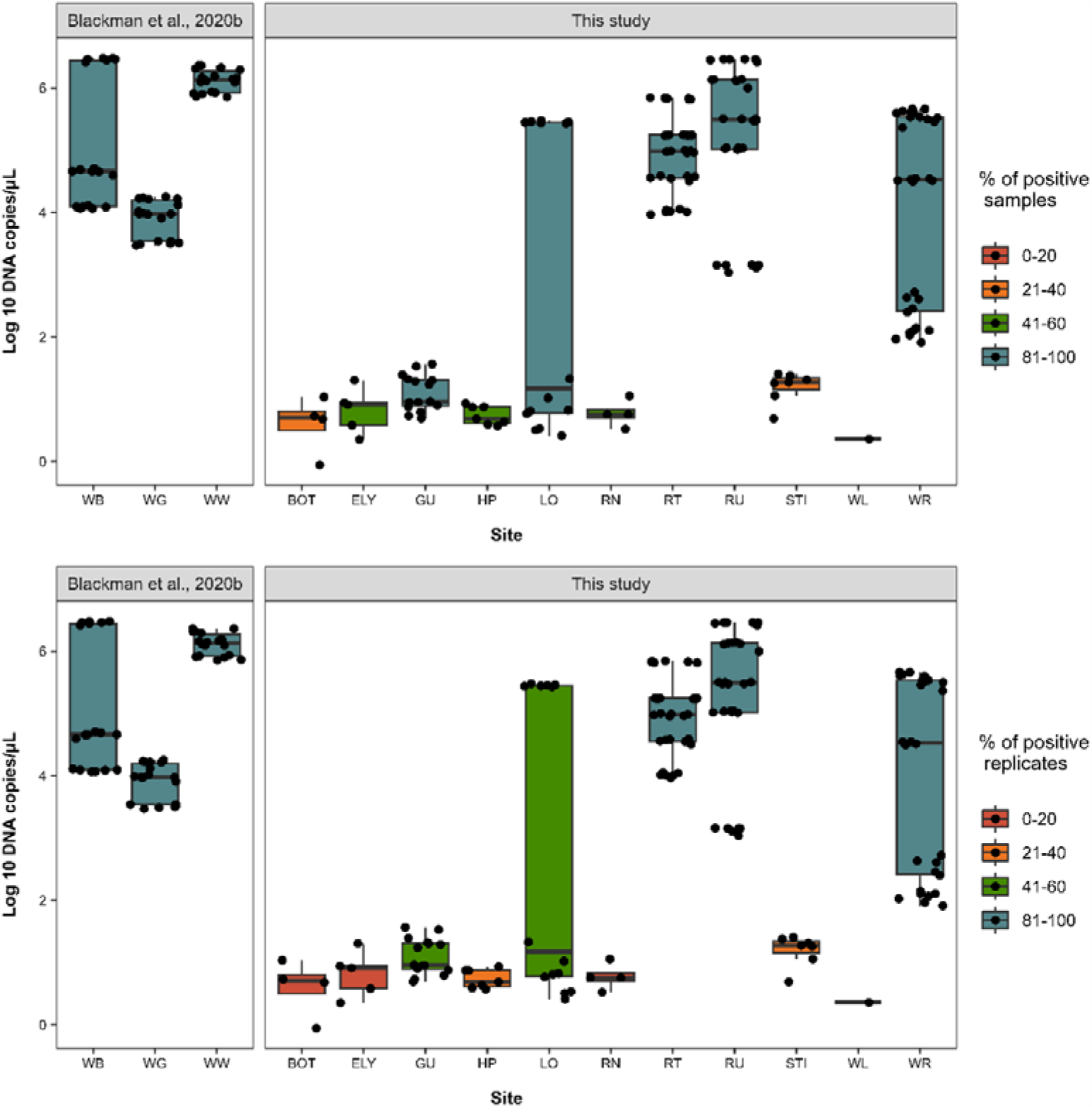
Log 10 DNA copies/μL and percentage of positive samples (top) and replicates (bottom) for samples from Blackman et al. (2020b) and from this study (total number of samples = 3 and 5, respectively; total number of replicates = 18 and 30, respectively). Only sites where quagga mussel amplification occurred are represented and site codes are the same as in Table 2 and Fig. 1. Each black dot represents a positive replicate. For visual purposes log 10 DNA copies/μL is being used instead of non-transformed data.

Regarding contamination, one negative field control (Rutland Water) showed amplification. However, contamination was only detected in one replicate (out of six) and at very low levels (Cq of 38.4) compared to the samples (Cq 19.9 – 31.4) and as such no data was discarded. All remaining field, extraction and PCR negative controls did not display any amplification.

## Discussion

In this study, the dDRB1 qPCR assay developed by Blackman et al. (2020b) was further developed to a probe-based assay (pDRB1) and successfully used to screen for quagga mussels in several locations in England, demonstrating its efficiency in detecting this invasive non-native species. The LOD for the pDRB1 assay was defined as 100 copies/μL and the DNA concentration obtained for the nine samples from Wraysbury River was significantly higher for the pDRB1 assay, demonstrating its increased sensitivity compared to the dDRB1 assay. Positive detections were obtained for 11 out of the 24 sites sampled in this study, with a stronger signal observed in locations where established populations were already known - Rutland Water, River Thames, and Walthamstow Reservoirs. Sites with low number of samples and/or replicates amplified as well as low copy numbers suggest a recent invasion and thus should be closely monitored to prevent population expansion. Our results demonstrate that quagga mussels are considerably more widespread in England than previously thought, with one of our positive sites – Holme Pierrepont water park – being the second furthest north they have been recorded so far. The detection of this species in a number of different rivers, recreational lakes and the canal system highlights the urgent need to implement biosecurity measures and limit further spread.

### pDRB1 assay

Thalinger et al. (2021) described a 5-level validation scale for targeted eDNA assays, ranging from “incomplete” (level 1) to “operational” (level 5). Initial testing was completed for the dDRB1 assay by Blackman et al. (2020b) through a series of mesocosm experiments and field trials, and the assay currently meets the criteria for level 3 (“essential”). In the present study, specificity and sensitivity of the new pDRB1 assay were further confirmed by testing in non-target and closely related species (zebra mussel, killer shrimp, demon shrimp, pea mussel, blue mussel, European oyster, common periwinkle and the common limpet), together with extensive field testing and determining the LOD. The additional development and testing of the DRB1 assay reported here brings the assay to level 4 (“substantial”) of the 5-level validation scale (Thalinger et al., 2021). By their definition, a positive detection at level 4 can be interpreted as the target being “very likely present”, whereas a negative detection means the target is likely to be absent, “assuming appropriate timing and replication in sampling”. To reach the highest level of validation, the influence of ecological and physical factors on eDNA availability and an estimation of detection probabilities from statistical modelling is needed, however this was beyond the scope of our study.

The nine samples previously collected and tested with the dDRB1 assay by Blackman et al. (2020b) and tested again here with the pDRB1 assay showed significantly different DNA concentrations. The probe-based assay yielded on average more than twice as much DNA copy numbers, demonstrating its increased sensitivity compared to the dye-based assay. Moreover, the LOD obtained for the pDRB1 assay in this study was 100 copies/μL. While Blackman et al. (2020b) had previously reported a LOD of 1E-04 ng/μL for the dDRB1 assay with SYBR green dye, and Marshall, Vanderploeg & Chaganti (2022) found the LOD to be 9.8E-05 ng/reaction for the same assay with the EvaGreen dye, the different units used to assess limits of detection (ng/μL and copies/μL) makes the direct comparison between LODs not fair.

### New records with eDNA

Quagga mussels are native to the Ponto-Caspian region in Eastern Europe, but they have been able to easily reach and colonise the rest of the continent in recent decades. They were first detected in GB in 2014 in the Wraysbury River (Aldridge et al., 2014) and they are now well established in the River Thames and its tributaries. Several locations within the Thames network are used to supply water to the Walthamstow Reservoirs, and quagga mussels are now established at this location too. Moreover, in 2020 they were found further north in the East Midlands in Rutland Water (Environment Agency, 2020). The detection rates for both samples and replicates at these three locations was 100%, and mean DNA copy number per site was in the order of 1.5E+05 to 9.1E+05 copies/μL, consistent with a high abundance of quagga mussels in these established locations.

Positive detections were obtained at an additional eight sites where quagga mussels were suspected and/or expected, but not previously confirmed. These sites are spread across four additional counties (Buckinghamshire, Cambridgeshire, Norfolk, Nottinghamshire) and include four rivers (the Cam, Great Ouse, Little Ouse and Nene), two popular water sports lakes (Holme Pierrepont and Willen Lake), and the Grand Union canal. This new distribution indicates that the species is now widespread and present in several major arteries as well as key recreational sites.

Of these eight new sites, detection rates were higher in the Grand Union Canal at Loughborough, and on the Little Ouse River at Southery, Norfolk. For these sites, positive detections were obtained in all five samples and in 47-50% of all replicates. The average DNA copy number at Southery was one order of magnitude lower than that of the three established populations, but still quite high compared to the remaining sites. The stretch of the Little Ouse River where the samples were collected is used for boat mooring, and a high number of boats were present at the time of sample collection. It is possible that mussels were attached to boats and/or mooring structures, which could have increased the concentration of quagga mussel eDNA at this location.

The remaining seven new sites displayed an average DNA copy number lower than the LOD defined for the pDRB1 assay. All field, extraction and PCR negative controls associated with these locations were negative, suggesting these were not false positives. According to Klymus et al. (2020), detections below the LOD can be expected to occur in eDNA studies due to the rarity of the target species, and should still be considered and not treated as noise. This could be an indication that the species is at an early point of invasion and population densities are not too high yet in these locations. Continuous monitoring at these sites is crucial and surveys at adjacent sites should be conducted frequently to prevent their further spread.

Despite the positive detections in several new locations, we were unable to detect quagga mussels in 13 sites out of the 24 sampled. According to the 5-level validation scale proposed by Thalinger et al. (2021), when the target species is not detected with a level 4 assay, it is likely that the target is absent “assuming appropriate timing and replication in sampling” has been conducted. The timing of sampling is unlikely to have caused any false negatives. Samples were collected during the summer period, whose temperatures favour breeding by quagga mussels. This has been shown to be associated with peaks in eDNA concentration in the environment (Spear et al., 2015). If quagga mussels were present at these sites, eDNA availability in the water would be expected to be high, thus increasing the likelihood of capturing eDNA during sample collection and their subsequent detection in the lab. Nevertheless, replication levels were not ideal for some of the sites. Due to lack of further access and safety concerns, samples were sometimes collected within a very small space (e.g. all samples were collected within 10 meters of each other at both Bishopbridge and Marham). If present, this could have prevented the collection and detection of quagga mussel eDNA at those sites, as eDNA is not homogeneously distributed in the water and collecting samples at different locations in the same site helps increase detection probabilities. Further surveys in more accessible areas and increased replication are needed at these locations to determine the population status of quagga mussels.

### Pathways

The extensive canal network in GB is a major potential pathway for the spread of quagga mussels. By attaching to boat hulls through their strong byssal threads, quagga mussels can easily reach and colonise new locations, and this is thought to be one of the main vectors for their spread (Karatayev & Burlakova, 2022). Moreover, canals are often connected to hotspots of human activity such as marinas, ports and industrial areas, facilitating further spread of INNS (Chapman et al., 2020). The detection of quagga mussel eDNA in samples from the Grand Union Canal in this study poses a serious concern. Due to its extension, being the longest canal in England (approximately 220 km long) connecting London and Birmingham, it could facilitate the further spread of this species in the southern part of the country. Recent lessons have been learnt from another Ponto-Caspian species, the demon shrimp (*D. haemobaphes*), which was first recorded in GB in 2012 and quickly became widespread once it entered the canal network (Johns et al., 2018).

Although to a lesser extent, rivers are also subject to human activities and as such are also potential routes that can be used by quagga mussels to spread further. Despite the previous belief that this species prefers canals and would not be found in fast flowing rivers (Aldridge et al., 2014), we were able to detect them in four new rivers - Cam, Great Ouse, Little Ouse and Nene - in addition to the river Thames. Even though the high flow rates of rivers when compared to canals might prevent quagga mussel veligers from settling in and establishing, they can still be used as pathways to reach new locations and as such the role of rivers on quagga mussel expansion cannot be overlooked.

Recreational activities such as water sports and fishing are also commonly associated with the spread of dreissenid mussels. Adults are able to strongly attach to hard structures such as boats, and veligers can be accidentally transported in nets and equipment, thus being introduced to new locations otherwise inaccessible to them. In addition to previously known reservoir sites, quagga mussels were detected at Holme Pierrepont water park in Nottingham and Willen Lake in Milton Keynes, both popular water sports facilities. The detection of quagga mussels in these locations highlights the urgent need for better biosecurity and “check, clean, dry” awareness campaigns at these sites, as it has been implemented at Grafham Water reservoir for example.

### Conclusions and future work

In the present study, the additional development and testing of the DRB1 assay brought the assay to level 4 of the 5-level validation scale defined by Thalinger et al. (2021), allowing to improve the readiness of the assay for routine monitoring and increasing the confidence in the results. At this level, a positive or negative eDNA result is a strong indication that the species is present or absent, respectively. Nevertheless, despite the improvements in the DRB1 assay, further work is needed before it can achieve the highest level (level 5) and be considered fully operational. This includes estimating detection probabilities via statistical modelling and investigating the influence of environmental and physical factors on eDNA detection.

Positive quagga mussel detections were obtained at 11 out of the 24 sites sampled, with a stronger signal being observed in locations where established populations were already known. Locations with lower eDNA concentrations could be at an early point of invasion and should be monitored closely to prevent further spread to adjacent water bodies. The 13 sites where quagga mussels were not detected should be considered high risk of invasion and therefore continued surveillance is needed, with the addition of more accessible areas and increased replication level in order to confirm if the species is absent or not in these locations.

Overall, our results demonstrate that quagga mussels are considerably more widespread in England than previously thought. Positive detections were obtained at eight new sites, spread across four counties, and including several rivers, recreational lakes and the canal system. Due to their high ecological and economic impacts, this highlights the urgent need to continuously monitor priority sites and potential pathways using eDNA and sensitive assays such as the pDRB1, in order to closely monitor their expansion. Information obtained from surveillance surveys can then be used by regulatory bodies and water companies to better assess the environmental impacts of risk activities (i.e. that can favour quagga mussel dissemination) such as water transfers between reservoirs or the authorization of fishing licences, ultimately improving site-specific biosecurity measures and minimising their further spread. Additionally, modelling environmental factors and hydrological connectivity together with eDNA data will help identify priority areas and thus focus monitoring and biosecurity efforts.

## CRediT authorship contribution

**Sara Peixoto:** conceptualization, data curation, formal analysis, investigation, methodology, visualization, writing – original draft preparation. **Rosetta C. Blackman:** methodology, writing – review & editing. **Jonathan Porter:** methodology, resources, validation, writing – review & editing. **Alan Wan:** methodology, resources, validation, writing – review & editing. **Chris Gerrard:** resources, writing – review & editing. **Ben Aston:** funding acquisition, writing – review & editing. **Lori Lawson Handley:** conceptualization, project administration, funding acquisition, supervision, writing – original draft preparation.

## Acknowledgements

We thank Gary Hodgetts, Julie Jackson, Rob Holland, Kylie Jones, Michael Drew, Richard Reynolds and Sam Westwood from Anglian Water and Abi Sheriden for their help with fieldwork and sample collection. We also thank Cuong Tang and Graham Sellers who provided lab advice, mainly regarding the extraction protocols. This work was supported by the Leeds-York-Hull Natural Environment Research Council (NERC) Doctoral Training Partnership (DTP) Panorama under grant NE/S007458/1 and by Yorkshire Water under WINEP project 7YW200058.

## References

Aldridge, D. C. (2023). Surveys for invasive species in Anglian Water reservoirs and water treatment works 2022. Cambridge Environmental Consulting.

Aldridge, D. C., Ho, S., & Froufe, E. (2014). The Ponto-Caspian quagga mussel, Dreissena rostriformis bugensis (Andrusov, 1897), invades Great Britain. Aquatic Invasions, 9(4), 529–535.

Bij de Vaate, A., & Beisel, J. N. (2011). Range expansion of the quagga mussel Dreissena rostriformis bugensis (Andrusov, 1897) in Western Europe: first observation from France. Aquatic Invasions, 6(Supplement 1), 71–74.

Blackman, R.C., Hänfling, B. & Lawson-Handley, L. (2018). The use of environmental DNA as an early warning tool in the detection of new freshwater invasive non-native species. CAB reviews, 13(010), 1–15.

Blackman, R., Bennucci, M., Donnelly, R., Hänfling, B., Harper, L. R., Sellers, G., & Lawson-Handley, L. (2020a). Simple, sensitive and species-specific assays for detecting quagga and zebra mussels (Dreissena rostriformis bugensis and D. polymorpha) using environmental DNA. Management of Biological Invasions, 11(2), 218–236.

Blackman, R. C., Ling, K. K. S., Harper, L. R., Shum, P., Hänfling, B., & Lawson□Handley, L. (2020b). Targeted and passive environmental DNA approaches outperform established methods for detection of quagga mussels, Dreissena rostriformis bugensis in flowing water. Ecology and Evolution, 10(23), 13248–13259.

Bustin, S. A., Benes, V., Garson, J. A., Hellemans, J., Huggett, J., Kubista, M., … & Wittwer, C. T. (2009). The MIQE guidelines: minimum information for publication of quantitative real-time PCR experiments. Clinical Chemistry, 55(4), 611–622.

Chakraborti, R. K., Madon, S., Kaur, J., & Gabel, D. (2013). Management and control of dreissenid mussels in water infrastructure facilities of the Southwestern United States. Quagga and zebra mussels. CRC Press, Boca Raton, 215–242.

Chakraborti, R. K., Madon, S., & Kaur, J. (2016). Costs for controlling dreissenid mussels affecting drinking water infrastructure: Case studies. American Water Works Association, 108(8), E442–E453.

Chapman, D. S., Gunn, I. D., Pringle, H. E., Siriwardena, G. M., Taylor, P., Thackeray, S. J., … & Carvalho, L. (2020). Invasion of freshwater ecosystems is promoted by network connectivity to hotspots of human activity. Global Ecology and Biogeography, 29(4), 645–655.

Connelly, N. A., O’Neill, C. R., Knuth, B. A., & Brown, T. L. (2007). Economic impacts of zebra mussels on drinking water treatment and electric power generation facilities. Environmental Management, 40, 105–112.

Di Muri, C., Handley, L. L., Bean, C. W., Li, J., Peirson, G., Sellers, G. S., … & Hänfling, B. (2020). Read counts from environmental DNA (eDNA) metabarcoding reflect fish abundance and biomass in drained ponds. bioRxiv, 2020–07.

Environment Agency. (2020, November 27). Quagga mussels found in the River Trent and Rutland Water. GOV.UK. https://www.gov.uk/government/news/quagga-mussels-found-in-the-river-trent-and-rutland-water

Feist, S. M., & Lance, R. F. (2021). Advanced molecular-based surveillance of quagga and zebra mussels: A review of environmental DNA/RNA (eDNA/eRNA) studies and considerations for future directions. NeoBiota, 66, 117–159.

Fonseca, V. G., Davison, P. I., Creach, V., Stone, D., Bass, D., & Tidbury, H. J. (2023). The application of eDNA for monitoring aquatic non-indigenous species: practical and policy considerations. Diversity, 15(5), 631.

Gallardo, B., & Aldridge, D. C. (2013). The ‘dirty dozen: socio-economic factors amplify the invasion potential of 12 highrisk aquatic invasive species in Great Britain and Ireland. Journal of Applied Ecology, 50(3), 757–766.

Gingera, T. D., Bajno, R., Docker, M. F., & Reist, J. D. (2017). Environmental DNA as a detection tool for zebra mussels Dreissena polymorpha (Pallas, 1771) at the forefront of an invasion event in Lake Winnipeg, Manitoba, Canada. Management of Biological Invasions, 8(3), 287.

Haltiner, L., Zhang, H., Anneville, O., De Ventura, L., Deweber, T., Hesselschwerdt, J., … & Dennis, S. (2022). The distribution and spread of quagga mussels in perialpine lakes north of the Alps. Aquatic invasions, 17(2), 153–173.

Johns, T., Smith, D. C., Homann, S., & England, J. A. (2018). Time-series analysis of a native and a non-native amphipod shrimp in two English rivers. BioInvasions Record, 7(2), 101–110.

Karatayev, A. Y., Burlakova, L. E., & Padilla, D. K. (1998). Physical factors that limit the distribution and abundance of Dreissena polymorpha (Pall.). Journal of Shellfish Research, 17(4), 1219–1235.

Karatayev, A. Y., Burlakova, L. E., & Padilla, D. K. (2015). Zebra versus quagga mussels: a review of their spread, population dynamics, and ecosystem impacts. Hydrobiologia, 746, 97–112.

Karatayev, A. Y., Karatayev, V. A., Burlakova, L. E., Mehler, K., Rowe, M. D., Elgin, A. K., & Nalepa, T. F. (2021). Lake morphometry determines Dreissena invasion dynamics. Biological Invasions, 23, 2489–2514.

Karatayev, A. Y., & Burlakova, L. E. (2022). What we know and don’t know about the invasive zebra (Dreissena polymorpha) and quagga (Dreissena rostriformis bugensis) mussels. Hydrobiologia, 1–74.

Klymus, K. E., Merkes, C. M., Allison, M. J., Goldberg, C. S., Helbing, C. C., Hunter, M. E., … & Richter, C. A. (2020). Reporting the limits of detection and quantification for environmental DNA assays. Environmental DNA, 2(3), 271–282.

Larson, J. H., Bailey, S. W., & Evans, M. A. (2022). Biofouling of a unionid mussel by dreissenid mussels in nearshore zones of the Great Lakes. Ecology and Evolution, 12(12), e9557.

Lawson Handley, L. (2015). How will the “molecular revolution” contribute to biological recording?. Biological Journal of the Linnean Society, 115(3), 750–766.

MacIsaac, H. J. (1996). Potential abiotic and biotic impacts of zebra mussels on the inland waters of North America. American Zoologist, 36(3), 287–299.

Marescaux, J., & Van Doninck, K. (2012). First records of Dreissena rostriformis bugensis (Andrusov, 1897) in the Meuse River. BioInvasions Record, 1(2).

Marshall, N. T., Vanderploeg, H. A., & Chaganti, S. R. (2022). Improving environmental DNA sensitivity for dreissenid mussels by targeting tandem repeat regions of the mitochondrial genome. Water, 14(13), 2069.

Mills, D. N., Chadwick, M., & Francis, R. A. (2019). Artificial substrate experiments to investigate potential impacts of invasive quagga mussel (Dreissena rostriformis bugensis, Bivalvia: Dreissenidae) on macroinvertebrate communities in a UK river. Aquatic Invasions, 14(2):365–383

Molloy, D. P., bij de Vaate, A., Wilke, T., & Giamberini, L. (2007). Discovery of Dreissena rostriformis bugensis (Andrusov 1897) in western Europe. Biological Invasions, 9, 871–874.

National Biodiversity Network Trust (2023). The National Biodiversity Network (NBN) Atlas. https://nbnatlas.org/.

Peñarrubia, L., Alcaraz, C., Vaate, A. B. D., Sanz, N., Pla, C., Vidal, O., & Viñas, J. (2016). Validated methodology for quantifying infestation levels of dreissenid mussels in environmental DNA (eDNA) samples. Scientific Reports, 6(1), 39067.

Prabhakaran, G. K., Sunkara, M., Raghavan, R., & Umapathy, G. (2023). Development of a species-specific qPCR assay for the detection of invasive African sharptooth catfish (Clarias gariepinus) using environmental DNA. Biological Invasions, 25(4), 975–982.

Quinn, A., Gallardo, B., & Aldridge, D. C. (2014). Quantifying the ecological niche overlap between two interacting invasive species: the zebra mussel (Dreissena polymorpha) and the quagga mussel (Dreissena rostriformis bugensis). Aquatic Conservation: Marine and Freshwater Ecosystems, 24(3), 324–337.

R Core Team (2022). R: A language and environment for statistical computing. R Foundation for Statistical Computing, Vienna, Austria. URL https://www.R-project.org/.

Roux, L. M. D., Giblot□Ducray, D., Bott, N. J., Wiltshire, K. H., Deveney, M. R., Westfall, K. M., & Abbott, C. L. (2020). Analytical validation and field testing of a specific qPCR assay for environmental DNA detection of invasive European green crab (Carcinus maenas). Environmental DNA, 2(3), 309–320.

Roy, H. E., Peyton, J., Aldridge, D. C., Bantock, T., Blackburn, T. M., Britton, R., … & Walker, K. J. (2014). Horizon scanning for invasive alien species with the potential to threaten biodiversity in Great Britain. Global Change Biology, 20(12), 3859–3871.

Salmaso, N., Ciutti, F., Cappelletti, C., Pindo, M., & Boscaini, A. (2022). First record of quagga mussel, Dreissena bugensis Andrusov, 1897, in Italy: morphological and genetic evidence in Lake Garda. BioInvasions Records, 11(4), 1031–1044.

Sanders, H., Mason, R. J., Mills, D. N., & Rice, S. P. (2022). Stabilization of fluvial bed sediments by invasive quagga mussels (Dreissena bugensis). Earth Surface Processes and Landforms, 47(14), 3259–3275.

Schloesser, D. W., Metcalfe-Smith, J. L., Kovalak, W. P., Longton, G. D., & Smithee, R. D. (2006). Extirpation of freshwater mussels (Bivalvia: Unionidae) following the invasion of dreissenid mussels in an interconnecting river of the Laurentian Great Lakes. The American Midland Naturalist, 155(2), 307–320.

Sellers, G. S., Di Muri, C., Gómez, A., & Hänfling, B. (2018). Mu-DNA: a modular universal DNA extraction method adaptable for a wide range of sample types. Metabarcoding and Metagenomics, 2, e24556.

Spear, S. F., Groves, J. D., Williams, L. A., & Waits, L. P. (2015). Using environmental DNA methods to improve detectability in a hellbender (Cryptobranchus alleganiensis) monitoring program. Biological Conservation, 183, 38–45.

Strayer, D. L., Adamovich, B. V., Adrian, R., Aldridge, D. C., Balogh, C., Burlakova, L. E., … & Jeschke, J. M. (2019). Long□term population dynamics of dreissenid mussels (Dreissena polymorpha and D. rostriformis): A cross□system analysis. Ecosphere, 10(4), e02701.

Thalinger, B., Deiner, K., Harper, L. R., Rees, H. C., Blackman, R. C., Sint, D., … & Bruce, K. (2021). A validation scale to determine the readiness of environmental DNA assays for routine species monitoring. Environmental DNA, 3(4), 823–836.

Tillotson, M. D., Kelly, R. P., Duda, J. J., Hoy, M., Kralj, J., & Quinn, T. P. (2018). Concentrations of environmental DNA (eDNA) reflect spawning salmon abundance at fine spatial and temporal scales. Biological Conservation, 220, 1–11.

Velde, G., & Platvoet, D. (2007). Quagga mussels Dreissena rostriformis bugensis (Andrusov, 1897) in the Main River (Germany). Aquatic Invasions, 2(3): 261–264.

Ward, J. M., & Ricciardi, A. (2007). Impacts of Dreissena invasions on benthic macroinvertebrate communities: a meta□analysis. Diversity and Distributions, 13(2), 155–165.

Wickham. H. (2016). ggplot2: elegant graphics for data analysis. Springer-Verlag New York.

Wright, D. A., Setzler-Hamilton, E. M., Magee, J. A., & Harvey, H. R. (1996). Laboratory culture of zebra (Dreissena polymorpha) and quagga (D. bugensis) mussel larvae using estuarine algae. Journal of Great Lakes Research, 22(1), 46–54.

