## supplementary material for "Mussels on the move: new records of the invasive non-native quagga mussel (*Dreissena rostriformis bugensis*) in Great Britain using eDNA and a new probe-based qPCR assay"

**DNA extraction of samples collected with Nature Metric filters (DNeasy Blood & Tissue Kit)**

1. Carefully wipe the outer surfaces of all filter units with 10% bleach and 70% ethanol and let them defrost.
2. Remove the plunger from a 3 ml luer lock syringe, as well as the top cap from the filter, and attach the syringe to the inlet of the filter.
3. Add 130 µl of 2 mg/ml of proteinase K to each filter. Make sure that the liquid is placed just above the filter.
4. Place the plunger back on the syringe and push the proteinase K into the filter. Carefully unscrew the syringe and cap the filter.
5. Incubate the capped filters at 56°C overnight.
6. Using a new 3 ml syringe expel the lysate into a 5 ml low retention microcentrifuge tube.
7. Add equal amounts of AL buffer to lysate in each tube, then vortex and incubate at 56°C for 30 minutes.
8. Add equal amounts of cold 100% ethanol as lysate to each tube and incubate for 10 minutes at room temperature.
9. For each sample transfer 600 μl of lysate to a spin column and centrifuge for 30 seconds at 6,000 x g. Discard flow-through and collection tube.
10. Repeat step 9 until all lysate is used. On the final spin centrifuge for 2 minutes.
11. Transfer spin column to new collection tube and add 500 μl of AW1 to spin column. Centrifuge for 1 minutes at 6,000 x g. Discard flow-through and collection tube.
12. Transfer spin column to new collection tube and add 500 μl of AW2 to spin column. Centrifuge for 3 minutes at 20,000 x g. Discard flow-through and collection tube.
13. Transfer spin column to a 1.5 low retention microcentrifuge tube, add 50 μl of AE buffer and incubate for 1 minute at room temperature. Centrifuge for 1 minute at 6,000 x g.
14. Add another 50 μl of AE buffer to the same tube and incubate for 1 minute at room temperature. Centrifuge for 1 minute at 6,000 x g.

**DNA extraction of samples collected with Sterivex filters (Mu-DNA protocol)**

**Reagents needed:** All reagents used in this protocol are from the Mu-DNA protocol (Sellers et al., 2018). Please refer to it for indications on how to make the solutions.

**Preparation:**

- Prepare 1.5 ml tubes with 300 μL Flocculant Solution and place in the fridge until required.
- Prepare 5 ml tubes with 2000 μL of Binding Solution and place in the oven at 55°C until required.
- Place Elution Buffer in the oven at 55°C until required.

**Protocol:**

1. Carefully wipe the outer surfaces of all filter units with 10% bleach and 70% ethanol and let them defrost.
2. Attach a 3 ml luer lock syringe to the inlet of the filter and push the Longmire’s buffer from the filter unit into a 2 ml sterile tube. Make sure to remove as much buffer as possible from the filter.
3. Spin the 2 ml tube with the buffer at 6,000 x g for 30 mins at room temperature. Discard the supernatant and dissolve the pellet (might not be visible) in 60 μL of Lysis Solution**.** Vortex for 15 s and transfer the liquid into the filter unit.
4. Keeping the outlet end closed, carefully add 660 μL of Lysis Solution, 180 μL of Tissue Lysis Additive (6% SDS) and 60 μL of 10 mg/ml proteinase K to the filter (using either a luer lock syringe or a 1000 µL pipet). Close the inlet, seal with parafilm and handshake vigorously for a few seconds. Do not vortex as this could dislodge the luer lock caps.
5. Incubate the samples overnight at 55°C on a rotating platform. The filter units should lay horizontally and be allowed to roll back and forth.
6. Handshake the filter unit vigorously. Attach a luer lock syringe to the inlet of the filter and pull (do not push) the lysate into the syringe. Transfer the lysate into a 2 ml tube and discard the filter.
7. Spin down for 15 s at 10,000 x g to remove any foam. Transfer as much liquid as possible (~900 μL) to the prepared tube with 300 μL (0.3x volume) of Flocculant Solution**.** Vortex to mix.
8. Incubate in the fridge for at least 10 min.
9. Centrifuge tubes at >10,000 x g for 1 min. Rotate the tubes by 180° (facing the opposite direction) in the centrifuge and centrifuge again for 1 min to make sure all the particles are pelleted.
10. Transfer 1000 μL of supernatant to the prepared tube with 2000 μL (2x volume) of Binding Solution, vortex to mix and centrifuge at >10,000 x g for 5 sec.
11. Transfer 670 μL of mixture to spin column and centrifuge at >10,000 x g for 10 sec, discard flow through. Repeat this until all mixture has passed through the spin column.
12. Add 500 μL of Wash Solution and centrifuge at >10,000 x g for 10 sec, discard flow through. Repeat this a second time.
13. Centrifuge spin columns at >10,000 x g for 2 min. Discard collection tube and contents.
14. Transfer spin column to a 1.5 mL labelled tube. Add 100 μL of Elution Buffer to spin column membrane, incubate at room temp for 1 min.
15. Centrifuge at >10,000 x g for 1 min.

**Table S1** Details of samples collected in this study, namely site name and sample code, coordinates, volume filtered and the TaqMan master mix used for both species-specific and inhibition qPCRs. Information on sample inhibition is provided in “Cq difference before” and “Cq difference after” columns, which indicate the difference between the average Cq of each sample and the no template reaction before and after 10x dilution, respectively.

| **Site name** | **Sample code** | **Latitude** | **Longitude** | **Volume filtered (mL)** | **TaqMan** | **Cq difference before** | **10X dilution** | **Cq difference after** |
| --- | --- | --- | --- | --- | --- | --- | --- | --- |
| Alton | ALT 1 | 51.979 | 1.157 | 720 | Universal | 7.557 | Yes | 0.471 |
| Alton | ALT 2 | 51.979 | 1.157 | 540 | Universal | 26.828 | Yes | 0.512 |
| Alton | ALT 3 | 51.979 | 1.158 | 660 | Universal | 26.828 | Yes | 0.149 |
| Alton | ALT 4 | 51.978 | 1.158 | 200 | Universal | 3.142 | Yes | 0.141 |
| Alton | ALT 5 | 51.978 | 1.157 | 420 | Universal | 0.501 | No | NA |
| Ardleigh | ARD 1 | 51.893 | 0.952 | 420 | Universal | 1.773 | No | NA |
| Ardleigh | ARD 2 | 51.893 | 0.952 | 480 | Universal | 1.701 | No | NA |
| Ardleigh | ARD 3 | 51.891 | 0.952 | 660 | Universal | 26.131 | Yes | 0.034 |
| Bedford WTW direct intake | BED 1 | 52.158 | -0.491 | 420 | Universal | 1.314 | No | NA |
| Bedford WTW direct intake | BED 2 | 52.158 | -0.491 | 420 | Universal | 0.163 | No | NA |
| Bedford WTW direct intake | BED 3 | 52.165 | -0.529 | 300 | Universal | 1.340 | No | NA |
| Bedford WTW direct intake | BED 4 | 52.165 | -0.526 | 360 | Universal | 3.191 | Yes | 0.057 |
| Bedford WTW direct intake | BED 5 | 52.165 | -0.526 | 420 | Universal | 0.368 | No | NA |
| Bishopbridge | BIS 1 | 53.406 | -0.450 | 300 | Universal | 0.664 | No | NA |
| Bishopbridge | BIS 2 | 53.406 | -0.450 | 300 | Universal | 0.910 | No | NA |
| Bishopbridge | BIS 3 | 53.406 | -0.450 | 300 | Universal | 8.093 | Yes | 0.095 |
| Bishopbridge | BIS 4 | 53.406 | -0.450 | 210 | Universal | 1.312 | No | NA |
| Bishopbridge | BIS 5 | 53.406 | -0.450 | 120 | Universal | 3.403 | Yes | 0.050 |
| Bottisham Lock | BOT 1 | 52.269 | 0.209 | 360 | Universal | 3.982 | Yes | 0.451 |
| Bottisham Lock | BOT 2 | 52.269 | 0.209 | 360 | Universal | 0.488 | No | NA |
| Bottisham Lock | BOT 3 | 52.269 | 0.209 | 360 | Universal | 2.527 | Yes | 0.032 |
| Bottisham Lock | BOT 4 | 52.269 | 0.209 | 360 | Universal | 0.269 | No | NA |
| Bottisham Lock | BOT 5 | 52.269 | 0.209 | 360 | Universal | 0.611 | No | NA |
| Costessey pits | COS 1 | 52.679 | 1.191 | 360 | Universal | 0.860 | No | NA |
| Costessey pits | COS 2 | 52.679 | 1.191 | 360 | Universal | 0.376 | No | NA |
| Costessey pits | COS 3 | 52.678 | 1.191 | 360 | Universal | 0.436 | No | NA |
| Costessey pits | COS 4 | 52.678 | 1.191 | 360 | Universal | 0.165 | No | NA |
| Costessey pits | COS 5 | 52.678 | 1.193 | 360 | Universal | 0.366 | No | NA |
| Covenham | COV 1 | 53.449 | 0.030 | 240 | Universal | 0.004 | No | NA |
| Covenham | COV 2 | 53.449 | 0.029 | 240 | Universal | 0.207 | No | NA |
| Covenham | COV 3 | 53.448 | 0.030 | 300 | Universal | 0.133 | No | NA |
| Covenham | COV 4 | 53.449 | 0.029 | 300 | Universal | 0.052 | No | NA |
| Covenham | COV 5 | 53.448 | 0.030 | 300 | Universal | 0.010 | No | NA |
| Elsham | ELS 1 | 53.513 | -0.492 | 115 | Universal | 26.828 | Yes | 0.692 |
| Elsham | ELS 2 | 53.513 | -0.492 | 44 | Universal | 2.597 | Yes | 0.152 |
| Elsham | ELS 3 | 53.513 | -0.492 | 102 | Universal | 26.828 | Yes | 0.338 |
| Elsham | ELS 4 | 53.513 | -0.492 | 33 | Universal | 8.608 | Yes | 0.098 |
| Elsham | ELS 5 | 53.513 | -0.492 | 109 | Universal | 3.412 | Yes | 0.030 |
| Ely waterfront/Marina | ELY 1 | 52.394 | 0.268 | 360 | Universal | 0.766 | No | NA |
| Ely waterfront/Marina | ELY 2 | 52.394 | 0.268 | 360 | Universal | 3.830 | Yes | 0.089 |
| Ely waterfront/Marina | ELY 3 | 52.394 | 0.268 | 360 | Universal | 0.484 | No | NA |
| Ely waterfront/Marina | ELY 4 | 52.394 | 0.268 | 360 | Universal | 0.422 | No | NA |
| Ely waterfront/Marina | ELY 5 | 52.395 | 0.269 | 360 | Universal | 0.003 | No | NA |
| Grafham | GRA 1 | 52.280 | -0.222 | 480 | Universal | 10.418 | Yes | 0.122 |
| Grafham | GRA 2 | 52.279 | -0.222 | 360 | Universal | 26.131 | Yes | 0.003 |
| Grafham | GRA 3 | 52.279 | -0.222 | 360 | Universal | 26.131 | Yes | 0.768 |
| Grafham | GRA 4 | 52.281 | -0.222 | 360 | Universal | 0.852 | No | NA |
| Grafham | GRA 5 | 52.278 | -0.222 | 360 | Universal | 5.703 | Yes | 0.128 |
| Grand Union canal | GU 1 | 52.784 | -1.218 | 1000 | Environmental | 0.030 | No | NA |
| Grand Union canal | GU 3 | 52.782 | -1.217 | 1000 | Environmental | 0.069 | No | NA |
| Grand Union canal | GU 5 | 52.780 | -1.216 | 900 | Environmental | 0.125 | No | NA |
| Grand Union canal | GU 7 | 52.778 | -1.215 | 1000 | Environmental | 0.120 | No | NA |
| Grand Union canal | GU 9 | 52.777 | -1.213 | 1000 | Environmental | 0.053 | No | NA |
| Holme Pierrepont park | HP 10 | 52.941 | -1.095 | 800 | Environmental | 0.013 | No | NA |
| Holme Pierrepont park | HP 2 | 52.951 | -1.071 | 1000 | Environmental | 0.154 | No | NA |
| Holme Pierrepont park | HP 4 | 52.952 | -1.077 | 1000 | Environmental | 0.176 | No | NA |
| Holme Pierrepont park | HP 6 | 52.948 | -1.082 | 600 | Environmental | 0.119 | No | NA |
| Holme Pierrepont park | HP 8 | 52.945 | -1.088 | 700 | Environmental | 0.046 | No | NA |
| Isleham Marina | ISL 1 | 52.355 | 0.420 | 360 | Universal | 1.158 | No | NA |
| Isleham Marina | ISL 2 | 52.355 | 0.420 | 360 | Universal | 0.596 | No | NA |
| Isleham Marina | ISL 3 | 52.354 | 0.420 | 360 | Universal | 0.129 | No | NA |
| Isleham Marina | ISL 4 | 52.354 | 0.420 | 360 | Universal | 3.401 | Yes | 0.411 |
| Isleham Marina | ISL 5 | 52.355 | 0.420 | 360 | Universal | 0.279 | No | NA |
| Little Ouse river | LO 1 | 52.500 | 0.367 | 1000 | Environmental | 0.035 | No | NA |
| Little Ouse river | LO 3 | 52.498 | 0.367 | 1000 | Environmental | 0.046 | No | NA |
| Little Ouse river | LO 5 | 52.495 | 0.368 | 1000 | Environmental | 0.113 | No | NA |
| Little Ouse river | LO 7 | 52.494 | 0.369 | 1000 | Environmental | 0.038 | No | NA |
| Little Ouse river | LO 9 | 52.492 | 0.371 | 1000 | Environmental | 0.051 | No | NA |
| Marham | MAR 1 | 52.678 | 0.548 | 360 | Universal | 0.188 | No | NA |
| Marham | MAR 2 | 52.678 | 0.549 | 360 | Universal | 0.027 | No | NA |
| Marham | MAR 3 | 52.679 | 0.549 | 360 | Universal | 0.398 | No | NA |
| Marham | MAR 4 | 52.679 | 0.549 | 360 | Universal | 0.098 | No | NA |
| Marham | MAR 5 | 52.678 | 0.548 | 360 | Universal | 0.162 | No | NA |
| River Nene | RN 1 | 52.566 | -0.221 | 1000 | Environmental | 0.047 | No | NA |
| River Nene | RN 3 | 52.566 | -0.224 | 1000 | Environmental | 0.113 | No | NA |
| River Nene | RN 5 | 52.566 | -0.227 | 1000 | Environmental | 0.074 | No | NA |
| River Nene | RN 7 | 52.566 | -0.231 | 1000 | Environmental | 0.064 | No | NA |
| River Nene | RN 9 | 52.567 | -0.235 | 1000 | Environmental | 0.099 | No | NA |
| River Thames | RT 1 | 51.563 | -0.709 | 1000 | Environmental | 0.135 | No | NA |
| River Thames | RT 3 | 51.565 | -0.712 | 1000 | Environmental | 0.026 | No | NA |
| River Thames | RT 5 | 51.567 | -0.713 | 1000 | Environmental | 0.032 | No | NA |
| River Thames | RT 7 | 51.569 | -0.712 | 1000 | Environmental | 0.021 | No | NA |
| River Thames | RT 9 | 51.570 | -0.712 | 800 | Environmental | 2.077 | No | NA |
| Rutland - Tinwell | RUT 1 | 52.642 | -0.500 | 300 | Universal | 1.341 | No | NA |
| Rutland - Tinwell | RUT 2 | 52.642 | -0.499 | 300 | Universal | 0.175 | No | NA |
| Rutland - Tinwell | RUT 3 | 52.642 | -0.497 | 300 | Universal | 0.905 | No | NA |
| Rutland - Tinwell | RUT 4 | 52.642 | -0.496 | 300 | Universal | 0.229 | No | NA |
| Rutland - Tinwell | RUT 5 | 52.643 | -0.494 | 300 | Universal | 0.440 | No | NA |
| Rutland Water | RU 1 | 52.644 | -0.655 | 1000 | Environmental | 0.082 | No | NA |
| Rutland Water | RU 3 | 52.642 | -0.649 | 1000 | Environmental | 0.129 | No | NA |
| Rutland Water | RU 5 | 52.640 | -0.645 | 900 | Environmental | 0.113 | No | NA |
| Rutland Water | RU 7 | 52.641 | -0.633 | 1000 | Environmental | 0.087 | No | NA |
| Rutland Water | RU 9 | 52.643 | -0.627 | 1000 | Environmental | 0.149 | No | NA |
| St Ives | STI 1 | 52.322 | -0.076 | 360 | Universal | 0.465 | No | NA |
| St Ives | STI 2 | 52.322 | -0.076 | 360 | Universal | 0.319 | No | NA |
| St Ives | STI 3 | 52.322 | -0.075 | 420 | Universal | 0.181 | No | NA |
| St Ives | STI 4 | 52.322 | -0.075 | 360 | Universal | 0.204 | No | NA |
| St Ives | STI 5 | 52.322 | -0.075 | 360 | Universal | 0.163 | No | NA |
| Staunton Harold reservoir | SH 1 | 52.815 | -1.442 | 1000 | Environmental | 0.012 | No | NA |
| Staunton Harold reservoir | SH 3 | 52.812 | -1.443 | 800 | Environmental | 0.182 | No | NA |
| Staunton Harold reservoir | SH 5 | 52.812 | -1.444 | 1000 | Environmental | 0.115 | No | NA |
| Staunton Harold reservoir | SH 7 | 52.811 | -1.446 | 500 | Environmental | 0.162 | No | NA |
| Staunton Harold reservoir | SH 9 | 52.811 | -1.447 | 1000 | Environmental | 0.186 | No | NA |
| Toft Newton Reservoir | TOF 1 | 53.373 | -0.448 | 145 | Universal | 1.049 | No | NA |
| Toft Newton Reservoir | TOF 2 | 53.374 | -0.448 | 110 | Universal | 0.865 | No | NA |
| Toft Newton Reservoir | TOF 3 | 53.374 | -0.448 | 165 | Universal | 0.465 | No | NA |
| Toft Newton Reservoir | TOF 4 | 53.373 | -0.449 | 175 | Universal | 0.332 | No | NA |
| Toft Newton Reservoir | TOF 5 | 53.372 | -0.448 | 88 | Universal | 0.089 | No | NA |
| Walthamstow reservoirs | WR 1 | 51.584 | -0.053 | 600 | Environmental | 0.091 | No | NA |
| Walthamstow reservoirs | WR 3 | 51.583 | -0.052 | 600 | Environmental | 0.113 | No | NA |
| Walthamstow reservoirs | WR 5 | 51.580 | -0.051 | 1000 | Environmental | 0.054 | No | NA |
| Walthamstow reservoirs | WR 7 | 51.582 | -0.049 | 1000 | Environmental | 0.095 | No | NA |
| Walthamstow reservoirs | WR 9 | 51.584 | -0.049 | 1000 | Environmental | 0.107 | No | NA |
| Willen lake | WL 1 | 52.054 | -0.723 | 600 | Environmental | 0.304 | No | NA |
| Willen lake | WL 3 | 52.055 | -0.715 | 600 | Environmental | 0.055 | No | NA |
| Willen lake | WL 5 | 52.053 | -0.715 | 500 | Environmental | 0.059 | No | NA |
| Willen lake | WL 7 | 52.051 | -0.714 | 400 | Environmental | 0.032 | No | NA |
| Willen lake | WL 9 | 52.048 | -0.715 | 500 | Environmental | 0.075 | No | NA |
| Wraysbury Gardens | WG_1 | 51.437 | -0.515 | 500 | Universal | 0.237 | No | NA |
| Wraysbury Gardens | WG_2 | 51.437 | -0.515 | 500 | Universal | 0.716 | No | NA |
| Wraysbury Gardens | WG_3 | 51.437 | -0.515 | 500 | Universal | 0.351 | No | NA |
| Wraysbury Bridge | WB_1 | 51.448 | -0.524 | 500 | Universal | 0.041 | No | NA |
| Wraysbury Bridge | WB_2 | 51.448 | -0.524 | 500 | Universal | 0.040 | No | NA |
| Wraysbury Bridge | WB_3 | 51.448 | -0.524 | 500 | Universal | 2.770 | Yes | 0.453 |
| Wraysbury Weir | WW_1 | 51.452 | -0.521 | 500 | Universal | 0.820 | No | NA |
| Wraysbury Weir | WW_2 | 51.452 | -0.521 | 500 | Universal | 0.711 | No | NA |
| Wraysbury Weir | WW_3 | 51.452 | -0.521 | 500 | Universal | 0.336 | No | NA |

**Table S2** Average Cq and average DNA copies/µL obtained for each sample (n=3) of each site (n=3) for both pDRB1 and dDRB1 assays.

| **Site name** | **Sample code** | **Average Cq** | **Average DNA copies/µL** | **Assay** |
| --- | --- | --- | --- | --- |
| Wraysbury Gardens | WG_1 | 25.165 | 7916.842 | dDRB1 |
| Wraysbury Gardens | WG_1 | 27.545 | 16334.305 | pDRB1 |
| Wraysbury Gardens | WG_2 | 26.166 | 4240.421 | dDRB1 |
| Wraysbury Gardens | WG_2 | 28.369 | 9420.177 | pDRB1 |
| Wraysbury Gardens | WG_3 | 27.929 | 1411.283 | dDRB1 |
| Wraysbury Gardens | WG_3 | 29.972 | 3234.392 | pDRB1 |
| Wraysbury Bridge | WB_1 | 22.378 | 20937.380 | dDRB1 |
| Wraysbury Bridge | WB_1 | 25.986 | 46097.418 | pDRB1 |
| Wraysbury Bridge | WB_2 | 24.393 | 5118.847 | dDRB1 |
| Wraysbury Bridge | WB_2 | 27.981 | 12169.778 | pDRB1 |
| Wraysbury Bridge | WB_3 | 18.644 | 462984.451 | dDRB1 |
| Wraysbury Bridge | WB_3 | 19.782 | 2869324.688 | pDRB1 |
| Wraysbury Weir | WW_1 | 16.057 | 1733209.249 | dDRB1 |
| Wraysbury Weir | WW_1 | 20.304 | 2041309.125 | pDRB1 |
| Wraysbury Weir | WW_2 | 17.116 | 827369.116 | dDRB1 |
| Wraysbury Weir | WW_2 | 20.918 | 1347928.375 | pDRB1 |
| Wraysbury Weir | WW_3 | 20.694 | 420085.720 | dDRB1 |
| Wraysbury Weir | WW_3 | 21.712 | 793622.250 | pDRB1 |

**Table S3** MIQE checklist (Bustin et al., 2009) for the newly developed pDRB1 assay.

| **Item to check** | **Importance** | **Details** |
| --- | --- | --- |
| Experimental design | | |
| Definition of experimental and control groups | E | 24 sites (including 1 canal, 2 lakes, 5 reservoirs and 16 rivers) spread throughout England and 3 sites on the River Wraysbury from Blackman et al. (2020b) |
| Number within each group | E | From this study: for 23 sites n = 5 and for 1 site n = 3  Samples from Blackman et al., (2020b), n = 3 |
| Assay carried out by the core or investigator's laboratory? | D | Investigator's lab |
| Acknowledgement of authors' contributions | D | Yes |
| Sample | | |
| Description | E | eDNA water samples |
| Volume/mass of sample processed | D | N/A |
| Microdissection or macrodissection | E | N/A |
| Processing procedure | E | Samples from this study: Water samples were filtered on site with either NatureMetrics or Sterivex filters. Water was pushed through the filter as many times as possible until the filter clogged, air was passed through the filter to dry it and 1 ml of Longmire’s buffer was added to each filter to preserve the sample.   Samples from Blackman et al. (2020b): 3 x 500 ml water samples were collected at each sampling point and vacuum filtered through sterile 47 mm diameter 0.45 μm cellulose nitrate membrane filters with pads (Whatman, GE Healthcare, UK) immediately after collection, using Nalgene filtration units (Thermo Fisher Scientific) in combination with a vacuum pump (15~20 in. Hg, Pall Corporation). |
| If frozen, how and how quickly? | E | N/A |
| If fixed, with what how quickly? | E | N/A |
| Sample storage conditions and duration (especially for FFPE samples) | E | Samples from this study: Filters were stored at room temperature while on fieldwork and stored at -20°C upon return to the lab (maximum 1 week).   Samples from Blackman et al. (2020b): All samples were stored in petri dishes at -20 °C until DNA extraction |
| Nucleic acid extraction | | |
| Procedure and/or instrumentation | E | Samples from this study: we used a modified version of the DNeasy Blood & Tissue Kit (Qiagen, UK) and a modified version of the Mu-DNA protocol (Sellers et al., 2018; Di Muri et al., 2020) for NatureMetrics and Sterivex filters, respectively. Both protocols are available in the supplementary material.   Samples from Blackman et al. (2020b): used a modified version of the Bolaski et al. (2008) protocol |
| Name of kit and details of any modifications | E |  |
| Source of additional reagents used | D | University of Hull |
| Details of DNase or RNAse treatment | E | N/A |
| Contamination assessment (DNA or RNA) | E | N/A |
| Nucleic acid quantification | E | Samples from this study: assessed using a Nanodrop 1000 spectrophotometer following the manufacturer's instructions.   Samples from Blackman et al. (2020b): assessed using a Qubit 2.0 following the manufacturer's instructions. |
| Instrument and method | E |  |
| Purity (A260/A280) | D |  |
| Yield | D |  |
| RNA integrity: method/instrument | E | N/A |
| RIN/RQI or Cq of 3' and 5' transcripts | E | N/A |
| Electrophoresis traces | D | N/A |
| Inhibition testing (Cq dilutions, spike, or other) | E | Samples were tested for inhibition using the Applied Biosystems TaqMan Exogenous Internal Positive Control Reagents (Fisher Scientific, UK). All eDNA samples were tested in duplicate and samples were considered to be inhibited if the average Cq of a sample was higher than the no template reaction by 2 or more cycles. Samples that showed inhibition were diluted 10x and re-run |
| Reverse Transcription | | |
| Complete reaction conditions | E | N/A |
| Amount of RNA and reaction volume | E | N/A |
| Priming oligonucleotide (if using GSP) and concentration | E | N/A |
| Reverse transcriptase and concentration | E | N/A |
| Temperature and time | E | N/A |
| Manufacturer of reagents and catalogue numbers | D | N/A |
| Cqs with and without reverse transcription | D | N/A |
| Storage conditions of cDNA | D | N/A |
| qPCR target information | | |
| Gene symbol | E | COI - cytochrome oxidase I |
| Sequence accession number | E | N/A |
| Location of amplicon | D | amplicon location: 196 – 384 |
| Amplicon length | E | 188 bp including primers |
| In silico specificity screen (BLAST, and so on) | E | N/A |
| Pseudogenes, retropseudogenes or other homologs? | D | N/A |
| Sequence alignment | D | N/A |
| Secondary structure analysis of amplicon | D | N/A |
| Location of each primer by exon or intron (if applicable) | E | N/A |
| What splice variants are targeted? | E | N/A |
| qPCR oligonucleotides | | |
| Primer sequences | E | forward (5’- 3’): GGA AAC TGG TTG GTC CCG AT reverse (5’- 3’): GGC CCT GAA TGC CCC ATA AT |
| RTPrimerDB Identification Number | D | N/A |
| Probe sequences | D | (5’- 3’): 6FAM - TCG GCG TTT AGT GAG GGC GGA TTT - QSY |
| Location and identity of any modifications | E | N/A |
| Manufacturer of oligonucleotides | D | IDT (primers) and ThermoFisher/Fisher Scientific (probe) |
| Purification method | D | HPLC |
| qPCR protocol | | |
| Complete reaction conditions | E | Volumes: 12.5 µL of TaqMan Environmental or Universal Master Mix 2.0, 1.6 µM of primers (forward and reverse combined), 0.05 µM of probe (Table 1), 0.64 mg/ml of BSA, 5.45 µL of water and 2 µL of sample.   qPCR program: 50°C for 10 min and 95°C for 10 min, followed by 45 cycles of 95°C for 15 seconds and 62°C for 1 min |
| Reaction volume and amount of cDNA/DNA | E | Reaction volume = 25 μL amount of DNA = 2 μL |
| Primer, (probe), Mg^2+^ and dNTP concentrations | E | 1.6 µM of primers (forward and reverse combined) and 0.05 µm of probe |
| Polymerase identity and concentration | E | TaqMan Environmental Master Mix 2.0 (Fisher Scientific, UK) TaqMan Universal Master Mix 2.0 (Fisher Scientific, UK) Both at a final concentration of 1x |
| Buffer/kit identity and manufacturer | E | N/A |
| Exact chemical composition of the buffer | D | N/A |
| Additives (SYBR Green I, DMSO, and so forth) | E | 0.64 mg/ml of BSA |
| Manufacturer of plates/tubes and catalog number | D | Applied Biosystems™ MicroAmp™ Fast Optical 96-Well Reaction Plate with Barcode, 0.1 mL (12142000, Fisher Scientific, UK) |
| Complete thermocycling parameters | E | 50°C for 10 min; 95°C for 10 min; 45 cycles of 95°C for 15 seconds and 62°C for 1 min |
| Reaction setup (manual/robotic) | D | Reactions were made manually |
| Manufacturer of qPCR instrument | E | StepOne-Plus™ Real-Time PCR system (Fisher Scientific/Thermo Fisher, UK) |
| qPCR validation | | |
| Evidence of optimisation (from gradients) | D | Three annealing temperatures (60, 62 and 63°C) were tested and all showed species-specific amplification of quagga mussel with no cross-amplification of the other species tested. Efficiency values were 102.9, 99.1 and 63.4% for the three annealing temperatures, respectively |
| Specificity (gel, sequence, melt, or digest) | E | N/A |
| For SYBR Green I, Cq of the NTC | E | N/A |
| Calibration curves with slope and y intercept | E | Slope range: -3.21 − -3.53; y-intercept range: 41.31−42.83 |
| PCR efficiency calculated from slope | E | 91.8 – 104.8% |
| CIs for PCR efficiency or SE | D | N/A |
| r2 of standard curve | E | 0.984-0.998 |
| Linear dynamic range | E | N/A |
| Cq variation at lower limit | E | The lowest standard (10 copies/µL) amplified at 93% of replicates |
| CIs throughout range | D | N/A |
| Evidence for LOD | E | The LOD, i.e. the lowest standard with at least 95% amplification, was 100 copies/µL |
| If multiplex, efficiency and LOD of each assay | E | N/A |
| Data analysis | | |
| qPCR analysis program (source, version) | E | StepOne Software version 2.3 |
| Method of Cq determination | E | Performed according to the default setting of the software above |
| Outlier identification and disposition | E | N/A |
| Results for NTCs | E | Eight wells of no-template negative control were included in all qPCR plates and showed no amplification |
| Justification of number and choice of reference genes | E | N/A |
| Description of normalisation method | E | We used standard curve methods |
| Number and concordance of biological replicates | D | From this study: for 23 sites n = 5 and for the remaining site n = 3 Samples from Blackman et al. (2020b), n = 3 |
| Number and stage (reverse transcription or qPCR) of technical replicates | E | Six qPCR replicates for each sample |
| Repeatability (intraassay variation) | E | N/A |
| Reproducibility (interassay variation, CV) | D | N/A |
| Power analysis | D | N/A |
| Statistical methods for result significance | E | N/A |
| Software (source, version) | E | N/A |
| Cq or raw data submission using RDML | D | N/A |

**Table S4** Cq values and DNA copies/µL obtained for all samples and replicates that showed amplification.

| **Site name** | **Sample code** | **Cq** | **DNA copies/µL** |
| --- | --- | --- | --- |
| Bottisham Lock | BOT 2 | 41.894 | 0.880 |
| Bottisham Lock | BOT 2 | 39.438 | 4.806 |
| Bottisham Lock | BOT 4 | 38.263 | 10.829 |
| Bottisham Lock | BOT 4 | 39.281 | 5.358 |
| Ely waterfront/Marina | ELY 1 | 38.938 | 8.752 |
| Ely waterfront/Marina | ELY 4 | 39.036 | 8.184 |
| Ely waterfront/Marina | ELY 4 | 37.721 | 20.222 |
| Ely waterfront/Marina | ELY 4 | 40.912 | 2.249 |
| Ely waterfront/Marina | ELY 5 | 40.139 | 3.830 |
| St Ives | STI 1 | 39.410 | 4.899 |
| St Ives | STI 1 | 37.340 | 20.501 |
| St Ives | STI 2 | 37.022 | 25.545 |
| St Ives | STI 2 | 38.191 | 11.383 |
| St Ives | STI 2 | 37.116 | 23.923 |
| St Ives | STI 2 | 37.507 | 18.259 |
| St Ives | STI 2 | 37.464 | 18.817 |
| Grand Union canal | GU 1 | 38.936 | 8.958 |
| Grand Union canal | GU 1 | 38.901 | 9.168 |
| Grand Union canal | GU 3 | 39.696 | 5.420 |
| Grand Union canal | GU 5 | 39.103 | 8.019 |
| Grand Union canal | GU 5 | 37.933 | 17.367 |
| Grand Union canal | GU 7 | 38.948 | 8.887 |
| Grand Union canal | GU 7 | 39.843 | 4.919 |
| Grand Union canal | GU 7 | 37.763 | 19.443 |
| Grand Union canal | GU 7 | 39.181 | 7.617 |
| Grand Union canal | GU 9 | 39.488 | 6.219 |
| Grand Union canal | GU 9 | 36.924 | 33.825 |
| Grand Union canal | GU 9 | 37.713 | 20.091 |
| Grand Union canal | GU 9 | 36.796 | 36.808 |
| Grand Union canal | GU 9 | 37.403 | 24.655 |
| Grand Union canal | GU 9 | 37.648 | 20.964 |
| Holme Pierrepont park | HP 10 | 39.271 | 4.884 |
| Holme Pierrepont park | HP 10 | 38.450 | 8.588 |
| Holme Pierrepont park | HP 4 | 38.653 | 7.468 |
| Holme Pierrepont park | HP 8 | 39.581 | 3.946 |
| Holme Pierrepont park | HP 8 | 39.664 | 3.728 |
| Holme Pierrepont park | HP 8 | 39.441 | 4.344 |
| Holme Pierrepont park | HP 8 | 38.645 | 7.511 |
| Little Ouse river | LO 1 | 38.775 | 5.892 |
| Little Ouse river | LO 1 | 38.596 | 6.674 |
| Little Ouse river | LO 3 | 39.561 | 3.401 |
| Little Ouse river | LO 5 | 23.417 | 270573.375 |
| Little Ouse river | LO 5 | 23.328 | 287887.188 |
| Little Ouse river | LO 5 | 23.353 | 282857.813 |
| Little Ouse river | LO 5 | 23.397 | 274389.750 |
| Little Ouse river | LO 5 | 23.372 | 279107.406 |
| Little Ouse river | LO 5 | 23.269 | 300035.063 |
| Little Ouse river | LO 7 | 39.943 | 2.604 |
| Little Ouse river | LO 9 | 36.928 | 21.428 |
| Little Ouse river | LO 9 | 38.644 | 6.457 |
| Little Ouse river | LO 9 | 39.643 | 3.212 |
| Little Ouse river | LO 9 | 37.953 | 10.467 |
| River Nene | RN 1 | 39.856 | 3.329 |
| River Nene | RN 5 | 38.088 | 11.349 |
| River Nene | RN 5 | 39.082 | 5.695 |
| River Nene | RN 9 | 39.078 | 5.714 |
| River Thames | RT 1 | 21.980 | 655017.688 |
| River Thames | RT 1 | 21.983 | 653662.438 |
| River Thames | RT 1 | 21.884 | 698016.250 |
| River Thames | RT 1 | 21.926 | 678640.500 |
| River Thames | RT 1 | 21.923 | 680257.500 |
| River Thames | RT 1 | 21.897 | 691911.750 |
| River Thames | RT 3 | 26.256 | 38878.109 |
| River Thames | RT 3 | 26.298 | 37812.781 |
| River Thames | RT 3 | 26.279 | 38279.215 |
| River Thames | RT 3 | 26.363 | 36213.402 |
| River Thames | RT 3 | 26.551 | 31988.484 |
| River Thames | RT 3 | 26.401 | 35326.191 |
| River Thames | RT 5 | 24.981 | 90228.523 |
| River Thames | RT 5 | 24.911 | 94509.758 |
| River Thames | RT 5 | 24.864 | 97508.852 |
| River Thames | RT 5 | 24.874 | 96831.609 |
| River Thames | RT 5 | 24.833 | 99522.734 |
| River Thames | RT 5 | 24.953 | 91915.500 |
| River Thames | RT 7 | 23.964 | 176681.609 |
| River Thames | RT 7 | 23.972 | 175673.672 |
| River Thames | RT 7 | 23.984 | 174388.281 |
| River Thames | RT 7 | 24.000 | 172454.875 |
| River Thames | RT 7 | 23.966 | 176422.688 |
| River Thames | RT 7 | 24.005 | 171957.594 |
| River Thames | RT 9 | 28.127 | 11293.130 |
| River Thames | RT 9 | 28.169 | 10985.740 |
| River Thames | RT 9 | 28.259 | 10351.801 |
| River Thames | RT 9 | 28.431 | 9240.614 |
| River Thames | RT 9 | 28.278 | 10226.050 |
| River Thames | RT 9 | 28.285 | 10173.094 |
| Rutland Water | RU 1 | 31.411 | 1086.496 |
| Rutland Water | RU 1 | 31.017 | 1424.257 |
| Rutland Water | RU 1 | 31.017 | 1424.446 |
| Rutland Water | RU 1 | 31.159 | 1292.223 |
| Rutland Water | RU 1 | 31.000 | 1441.373 |
| Rutland Water | RU 1 | 31.020 | 1421.807 |
| Rutland Water | RU 3 | 19.950 | 2875904.000 |
| Rutland Water | RU 3 | 19.939 | 2898461.750 |
| Rutland Water | RU 3 | 19.943 | 2889469.750 |
| Rutland Water | RU 3 | 20.089 | 2614143.000 |
| Rutland Water | RU 3 | 19.947 | 2882463.500 |
| Rutland Water | RU 3 | 19.984 | 2808869.750 |
| Rutland Water | RU 5 | 24.784 | 103570.047 |
| Rutland Water | RU 5 | 24.714 | 108648.539 |
| Rutland Water | RU 5 | 24.727 | 107699.516 |
| Rutland Water | RU 5 | 24.798 | 102568.609 |
| Rutland Water | RU 5 | 24.754 | 105707.938 |
| Rutland Water | RU 5 | 24.707 | 109173.500 |
| Rutland Water | RU 7 | 21.105 | 1299377.625 |
| Rutland Water | RU 7 | 21.081 | 1321098.750 |
| Rutland Water | RU 7 | 21.024 | 1374016.000 |
| Rutland Water | RU 7 | 21.487 | 999127.375 |
| Rutland Water | RU 7 | 21.043 | 1355820.625 |
| Rutland Water | RU 7 | 21.034 | 1364297.375 |
| Rutland Water | RU 9 | 23.153 | 317896.813 |
| Rutland Water | RU 9 | 23.140 | 320734.656 |
| Rutland Water | RU 9 | 23.236 | 300169.625 |
| Rutland Water | RU 9 | 23.242 | 298885.406 |
| Rutland Water | RU 9 | 23.147 | 319043.000 |
| Rutland Water | RU 9 | 23.237 | 300093.281 |
| Walthamstow reservoirs | WR 1 | 33.537 | 250.721 |
| Walthamstow reservoirs | WR 1 | 32.760 | 430.098 |
| Walthamstow reservoirs | WR 1 | 34.642 | 116.378 |
| Walthamstow reservoirs | WR 1 | 33.351 | 285.365 |
| Walthamstow reservoirs | WR 1 | 32.841 | 406.596 |
| Walthamstow reservoirs | WR 1 | 32.475 | 524.356 |
| Walthamstow reservoirs | WR 3 | 34.510 | 127.594 |
| Walthamstow reservoirs | WR 3 | 34.966 | 92.902 |
| Walthamstow reservoirs | WR 3 | 34.389 | 138.765 |
| Walthamstow reservoirs | WR 3 | 35.154 | 81.566 |
| Walthamstow reservoirs | WR 3 | 34.766 | 106.738 |
| Walthamstow reservoirs | WR 3 | 34.494 | 128.996 |
| Walthamstow reservoirs | WR 5 | 22.940 | 395061.063 |
| Walthamstow reservoirs | WR 5 | 22.889 | 409266.094 |
| Walthamstow reservoirs | WR 5 | 22.732 | 456379.750 |
| Walthamstow reservoirs | WR 5 | 22.967 | 387471.094 |
| Walthamstow reservoirs | WR 5 | 22.840 | 423232.000 |
| Walthamstow reservoirs | WR 5 | 22.728 | 457459.281 |
| Walthamstow reservoirs | WR 7 | 23.398 | 287246.188 |
| Walthamstow reservoirs | WR 7 | 23.258 | 316678.719 |
| Walthamstow reservoirs | WR 7 | 23.151 | 340970.719 |
| Walthamstow reservoirs | WR 7 | 23.714 | 230634.797 |
| Walthamstow reservoirs | WR 7 | 23.189 | 332248.125 |
| Walthamstow reservoirs | WR 7 | 23.153 | 340631.563 |
| Walthamstow reservoirs | WR 9 | 26.580 | 31492.461 |
| Walthamstow reservoirs | WR 9 | 26.414 | 35344.160 |
| Walthamstow reservoirs | WR 9 | 26.529 | 32623.395 |
| Walthamstow reservoirs | WR 9 | 26.518 | 32884.672 |
| Walthamstow reservoirs | WR 9 | 26.462 | 34197.934 |
| Walthamstow reservoirs | WR 9 | 26.425 | 35090.160 |
| Willen lake | WL 3 | 40.399 | 2.285 |
